# A graph neural network model to estimate cell-wise metabolic flux using single cell RNA-seq data

**DOI:** 10.1101/2020.09.23.310656

**Authors:** Norah Alghamdi, Wennan Chang, Pengtao Dang, Xiaoyu Lu, Changlin Wan, Silpa Gampala, Zhi Huang, Jiashi Wang, Qin Ma, Yong Zang, Melissa Fishel, Sha Cao, Chi Zhang

## Abstract

The metabolic heterogeneity, and metabolic interplay between cells and their microenvironment have been known as significant contributors to disease treatment resistance. However, with the lack of a mature high-throughput single cell metabolomics technology, we are yet to establish systematic understanding of intra-tissue metabolic heterogeneity and cooperation phenomena among cell populations. To mitigate this knowledge gap, we developed a novel computational method, namely scFEA (single cell Flux Estimation Analysis), to infer single cell fluxome from single cell RNA-sequencing (scRNA-seq) data. scFEA is empowered by a comprehensively reconstructed human metabolic map into a factor graph, a novel probabilistic model to leverage the flux balance constraints on scRNA-seq data, and a novel graph neural network based optimization solver. The intricate information cascade from transcriptome to metabolome was captured using multi-layer neural networks to fully capitulate the non-linear dependency between enzymatic gene expressions and reaction rates. We experimentally validated scFEA by generating an scRNA-seq dataset with matched metabolomics data on cells of perturbed oxygen and genetic conditions. Application of scFEA on this dataset demonstrated the consistency between predicted flux and metabolic imbalance with the observed variation of metabolite abundance in the matched metabolomics data. We also applied scFEA on five publicly available scRNA-seq and spatial transcriptomics datasets and identified context and cell group specific metabolic variations. The cell-wise fluxome predicted by scFEA empowers a series of downstream analysis including identification of metabolic modules or cell groups that share common metabolic variations, sensitivity evaluation of enzymes with regards to their impact on the whole metabolic flux, and inference of cell-tissue and cell-cell metabolic communications.

## INTRODUCTION

Metabolic dysregulation is a hallmark of many disease types including cancer, diabetes, cardiovascular disease and Alzheimer’s disease (Mattson and Chan 2001; Rask et al. 2001; Matsuzawa 2006; Dunn et al. 2014; Hirschey et al. 2015; Kochanek et al. 2019; Sun et al. 2020a). In cancer, the diseased cells are well understood to rewire their metabolism and energy production to support rapid proliferation, sustain viability, and promote acquired drug resistance (Thompson et al. 2005; DeBerardinis et al. 2008; Hanahan and Weinberg 2011; Ward and Thompson 2012). Here, the diseased cells often react differently to the microenvironmental stress. Such heterogeneity often results in an increased repertoire of possible cellular responses to compromise the efficacy of drug therapies, and synergistic cooperation among the cells that can ultimately enhance the survival of the entire population (Bishop et al. 2007; Lidstrom and Konopka 2010). The metabolome is an excellent indicator of phenotypic heterogeneity due to its high dynamics and plasticity (Zenobi 2013): one may expect to see a subset of cancerous cells, such as circulating tumor cells, that display abnormally high metabolic rates compared with many others with normal metabolism, and rare cells that successfully cope with microenvironmental stress, whereas the others die. Unfortunately, current high-throughput metabolic profiling has been largely applied to bulk cell or tissue samples, from which we could only observe an averaged metabolic signal over a large number of cells, while single cell metabolomics is still in its infancy, due by its relatively low throughput and low sensitivity (Zenobi 2013; Fessenden 2016; Emara et al. 2017; Zampieri et al. 2017; Ali et al. 2019; Duncan et al. 2019; Ahl et al. 2020). Hence, our understanding of metabolic dysregulation of human disease has been immensely limited by our technology to study the metabolic landscape at single-cell level and in the context of their tissue microenvironment (Jaenisch and Bird 2003; Feinberg 2007; Heintzman et al. 2007; Harris et al. 2010; Dunham et al. 2012; Roadmap Epigenomics et al. 2015; Robertson-Tessi et al. 2015; Kim and DeBerardinis 2019).

Single cell RNA-Seq (scRNA-seq) data has been widely utilized to characterize cell type specific transcriptional states in a complex tissue. Large amount of scRNA-seq data carry the potential to deliver information on a cell’s functioning state and its underlying phenotypic switches (Vasdekis and Stephanopoulos 2015; Damiani et al. 2019a; Evers et al. 2019a; Honkoop et al. 2019; Saurty-Seerunghen et al. 2019; Xiao et al. 2019a; Levine et al. 2020; Rohlenova et al. 2020; Xiao et al. 2020; Zhang et al. 2020). Realizing the strong connections between transcriptomic and metabolomic profiles (Hirayama et al. 2009; Lee et al. 2012; Mehrmohamadi et al. 2014; Damiani et al. 2019b; Xiao et al. 2019b; Wagner et al. 2020), scRNA-Seq has found its application in portraying metabolic variations. Most of the existing studies examined single cell metabolic changes relying on differential expression and enrichment analysis of key metabolic enzymes and pathways (Vasdekis and Stephanopoulos 2015; Evers et al. 2019a; Honkoop et al. 2019; Saurty-Seerunghen et al. 2019; Xiao et al. 2019a; Levine et al. 2020; Rohlenova et al. 2020; Xiao et al. 2020), without considering individual metabolite nodes in a metabolic pathway, or the mass balance constraints of metabolic network. Studies coupling single cell transcriptomics data and the Flux Balance Analysis (FBA) at steady-state framework have only recently emerged (Damiani et al. 2019a; Zhang et al. 2020). The FBA describes the potential flux over the topological structure of a metabolic network, with a set of equations governing the mass balance at steady state. The advantage of incorporating FBA is two-fold: considering the chemical stoichiometry in FBA could lead to more accurate estimation of the metabolite abundance; flux estimation for each individual metabolite can be solved, leading to high-resolution characterization of the metabolic profiling. Damiani et al developed scFBA that utilizes the cell group specific gene expression status derived from scRNA-seq data to regularize the network topology for FBA (Damiani et al. 2019a). Wagner et al proposed a method, namely Compass, which maximizes the coherence between scRNA-seq expression profile and predicted flux in solution space of FBA (Wagner et al. 2020). Compass characterizes the cell heterogeneity within a genome-scale metabolic model and utilizes neighboring cells to increase the robustness of single cell FBA. However, as stated in the original works, the stringent flux balance and steady-state assumption in scFBA and Compass may undermine the rationality of the optimization goal in the case of cancer, because the cancer cells and other cell types are constantly facing severe “imbalance” for many metabolites, driven by the substantial level of metabolic stress within the tumor microenvironment. Another limitation of the FBA-based methods is that the single cells’ gene expression is not used directly to model metabolic flux. Both scFBA and Compass used single cell gene expression as continuous or discrete constraint to guide the search of an optimal solution in the solution space of flux balance condition, which ignored the non-linear relationship between gene expression and reaction rates governed by Michaelis-Menten kinetic model. In addition, it is noteworthy that these models are intended for modeling the fluxes for cells of pre-defined groups, instead of at a single cell resolution, and they are restricted to a small portion of the whole metabolic map. In summary, there is a lack of methodology to predict the single cell metabolic profile using scRNA-seq data, that could flexibly incorporate the flux balance constraint and model the non-linear relationship between the metabolome and the transcriptome. Therefore, to unravel the principles of how malignant transformation affects the metabolic phenotypes of the heterogeneous cell types within the tumor microenvironment, it is urgent to design advanced computational tools to empower a reliable estimation of cell-wise metabolic flux and states by designing more appropriate and sophisticated model to translate single cell transcriptomes to single cell fluxomes (Damiani et al. 2019b; Evers et al. 2019b).

Computational challenges to estimate cell-wise metabolic flux arise from the following aspects: (1) multiple key factors determine cells’ metabolic states, including exogeneous nutrient availability in the tissue microenvironment, leading to a disjunction of cell type specific markers and metabolic phenotypes, and making conventional single cell clustering methods inapplicable; (2) the whole metabolic network is of high complexity, hence a proper computational reduction and reconstruction of the network is needed to reach a balance between resolution of metabolic state characterization and computational feasibility; (3) the intricate non-linear dependency between transcripts level and metabolic reaction rates calls for a more sophisticated model to fully capitulate the relationships; and (4) alternations on different enzymes of a metabolic pathway may result in common metabolic phenotypes, however, exactly which enzymes share such common effect to the metabolic flux change remains largely unknown.

In this study, we developed a novel computational method, namely **<u>s</u>**ingle-**<u>c</u>**ell **<u>F</u>**lux **<u>E</u>**stimation **<u>A</u>**nalysis (scFEA), to estimate the relative rate of metabolic flux at single cell resolution from scRNA-Seq data. Specifically, scFEA is empowered by the following computational innovations that can effectively solve the above challenges: (i) an optimization function derived based upon a probabilistic model to leverage the flux balance constraints among a large number of single cells with possibly varied metabolic fluxomes, (ii) a metabolic map reduction approach based on network topology and gene expression status, (iii) a multi-layer neural network model to capture the non-linear dependency of metabolic flux on the enzymatic gene expressions, and (iv) a novel graph neural network architecture and solution to maximize the overall flux balance of intermediate substrates through all cells. The central hypotheses of scFEA are (1) the flux variations of a metabolic module, composed by closely connected reactions, can be modeled as a non-linear function of the transcriptomic-level changes of the catalyzing enzymes and (2) the total flux imbalance of all intermediate substrates should be minimized through all single cell samples. The cell-wise fluxome estimated by scFEA enables a series of downstream analysis including identification of cell or tissue level metabolic stress, sensitivity evaluation of enzymes to the metabolic rewiring, and inference of cell-tissue and cell-cell metabolic exchanges. To validate scFEA, we generated an scRNA-seq dataset with matched tissue level metabolomic profiles under different biochemical perturbations. Applications of scFEA on synthetic datasets, the newly generated dataset with matched scRNA-Seq and metabolic profiles, and six other independent real-world datasets, validated the prediction accuracy, robustness, and biological interpretability of scFEA.

## RESULTS

### Systems biology considerations, hypotheses, and analysis pipeline of scFEA

The reaction rate of a simple enzyme catalyzed metabolic reaction follows the Michaelis-Menten model: 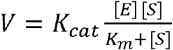, which is a non-linear function of enzyme concentration [*E*], substrate concentration [*S*], and kinetic parameters *K*_cat_ and *K*_*m*_. However, existing biotechnology does not enable a simultaneous high throughput measurement of these hyper-parameters, nor the metabolic flux from single cell samples. In sight of this limitation, we provide a computational estimation of cell-wise fluxome from a different angle. Based on the Michaelis-Menten model, the metabolic rate of a reaction depends more on the concentration of enzymes than substrates when the substrate concentration is close to saturation, i.e., when 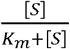 is large. In addition, the flux balance of intermediate metabolites largely constrains the fluxome distribution of connected metabolic reactions. Hence, the flux of a reaction chain is mostly determined by the enzyme variation of the rate limiting steps. However, such rate limiting steps are always unknow because the flux distribution, substrate concentration, and kinetic parameters highly depend on the physiological and biochemical conditions of the cells. Based on these considerations, we developed scFEA, to estimate cell-wise metabolic flux from scRNA-seq data. scFEA consists of three major computational components, namely (1) network reduction and reconstruction, (2) estimation of cell-wise metabolic flux by a graph neural network based approach, and (3) downstream analyses including estimation of metabolic stress, perturbation of metabolic genes, and clustering of cells with different metabolic states (**Figure 1**). The required input of scFEA is an scRNA-seq dataset, while optional inputs, including cell group labels or subset of metabolic reactions of interest, can be specified for additional analysis.

**Figure 1.**
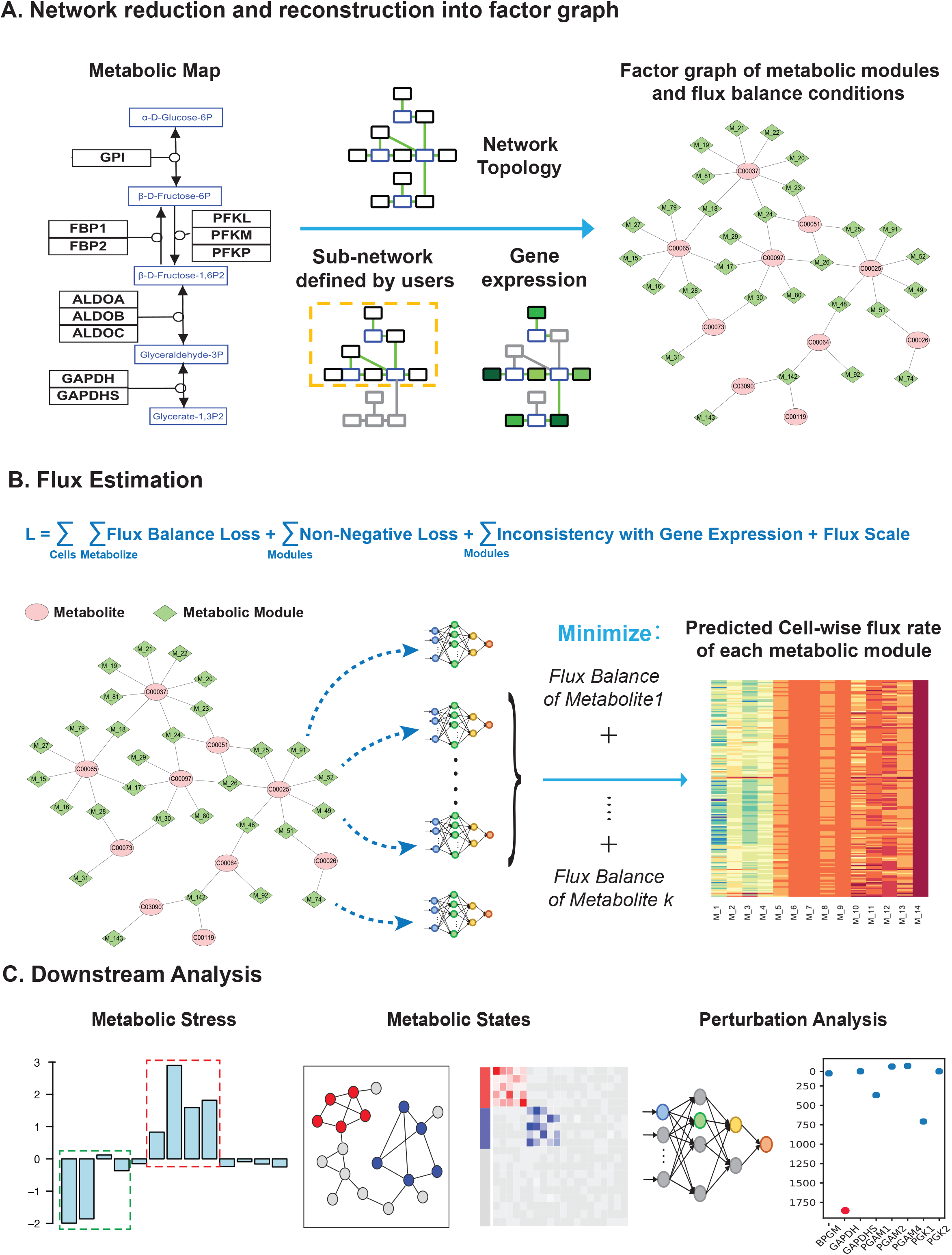
The computational framework of scFEA. (A) Metabolic reduction and reconstruction. A metabolic map was reduced and reconstructed into a factor graph based on network topology, significantly non-zero gene expressions and users’ input. (B) A novel graph neural network architecture based prediction of cell-wise fluxome. A loss function (L) composed by loss terms of flux balance, non-negative flux, coherence between predicted flux and gene expression, and constraint of flux scale were utilized to estimate cell-wise metabolic flux from scRNA-seq data. See detailed models and formulations in Results and Methods. (C) Downstream analysis of scFEA is provided, including inference of metabolic stress, cell and module clusters of distinct metabolic states, and the genes of top impact to the whole metabolic flux.

To reduce the complexity of the metabolic map, we reconstructed it into a factor graph composed by connected metabolic modules as variables and intermediate metabolites as factors (**Figure 1A**). Specifically, connected reactions are merged into one module if changes in their reaction rates do not affect the rates of the other reactions conditional to a fixed flux rate of the module. In other words, the solution we obtained for estimating the flux of a module stays the same with or without merging the reactions, under the flux balance condition. This approach increases the robustness of flux estimation and reduces the computational complexity.

The central computational component of scFEA is a novel graph neural network architecture, which models cell-wise metabolic flux of each metabolic module by using gene expression levels of the catalyzing enzymes in each individual cell (**Figure 1B**). We hypothesize that the metabolic flux throughout all the single cells collected from a tissue sample should minimize the overall imbalance of the in-/out-flux of intermediate substrates, i.e., maintaining the maximal flux balance of intermediate substrates through all the cells. The rationality of this assumption is that cells within the same tissue inevitably exchange the metabolites, and hence the total flux balance of intermediate substrates throughout all the single cells collected from one tissue sample are more robust than in individual cells. In scFEA, we utilize the variations in gene expression to reflect the protein level change of enzymes and transporters. This hypothesis can also be supported by many existing studies that reveal the high explainability of the transcriptome for the protein level of enzymes (Schnell 2014; Roadmap Epigenomics et al. 2015; Liu et al. 2016). We assume the flux variations of a connected metabolic module generally impacts its neighboring modules, which can be reflected by aggregating the expression variations of the genes in its neighborhood over the metabolic network. The non-linear dependency between gene expression change and metabolic flux of each module is modeled as a fully connected neural network of 2-4 layers, which could be considered as a non-linear approximation of the Michaelis-Menten model. To solve the neural network parameters, scFEA minimizes a loss function that mimics the overall flux imbalance of all modules in all cells, with further non-negativity and other prior assumptions on the module fluxome. The large single cell sample size in scRNA-seq data grants sufficient statistical power to detect the flux variations and avoids the overfitting of the neural network training (see details in methods). It is noteworthy the parameters of the neural network of each module describe the impact of each gene’s expression to the predicted flux, which measures the sensitivity of the metabolic balance to the variations of the genes. Genes with higher impact indicate rate limiting reactions under the particular context.

The estimated cell-wise metabolic flux enables the prediction of (i) the metabolites or pathways with high imbalance in certain cell group, (ii) groups of metabolic modules or cells with varied metabolic states, and (iii) the metabolic genes whose perturbation highly impacts the overall metabolic flux (**Figure 1C**). In this study, we mainly focus on solving cell-wise metabolic flux and states, and method validations in human cells. A capability for mouse data analysis is also provided in the software package of “scFEA”.

### Metabolic map reduction and reconstruction

The whole metabolic network in human and mouse have been well studied. However, while databases including the Kyoto Encyclopedia of Genes and Genomes (KEGG) provide well categorized metabolic pathways and the comprehensive set of metabolic genes (Kanehisa and Goto 2000), the network topological structure needs to be further optimized for fluxome estimation, due to the following reasons: (1) the flux balance relationships among different reactions could vary depending on the optimization goal or computational assumption, such as flux balance condition of carbon, redox or pH, (2) the network complexity needs to be reduced to enable computational feasibility, and (3) a manual correction and annotation of the directions of reactions and transporters is in need. In addition, cells of different types or physiological states naturally have varied metabolic states. In scFEA, we first manually curated and annotated the metabolic map of human and mouse retrieved from KEGG database. The global metabolic map of human and mouse is further reduced and reconstructed into a factor graph based on its topological property. ScFEA also enables the selection of a connected sub-network in the global metabolic network for flux estimation.

#### Collection of human and mouse metabolic map

The metabolic map consists of pathways and reactions that fall under four major types, namely import, metabolism, biosynthesis, and export. To ensure a comprehensive coverage of the global metabolic map, we collected reactions of metabolism and biosynthesis as well as transporters for import and export from different data sources. Specifically, metabolic reactions were directly retrieved from metabolic pathways in KEGG database (Kanehisa and Goto 2000); the transporters and annotations of import and export reactions were accessed from the transporter classification database (Saier et al. 2006); biosynthesis reactions were collected from the biosynthesis pathways encoded in KEGG and curated by using additional literatures (see details in Supplementary Methods). The final metabolic map covers the metabolism, transport, and biosynthesis of carbohydrate, amino acids, fatty acids and lipids, glycan, and nucleic acids in human and mouse, including 862 genes of 390 enzymes, 1880 reactions, 1219 metabolites, and 116 transporter genes of 35 metabolites in human. Completes gene and reaction lists of the collected human metabolic map is given in Supplementary Table S1.

#### Reconstruction of metabolic map into a factor graph

The metabolic reaction naturally forms a directed factor graph when considering the reaction as a variable and each metabolite as a factor. A directed factor graph was first reconstructed by the stoichiometric matrix of all reactions in the global metabolic map, in which variable, factor, and directed edge are reactions, metabolites, and whether or not a reaction involves a metabolite as the substrate or product, respectively. In this study, we use a flux balance assumption of carbon-based metabolites, and hence 273 compounds that do not affect the flux balance of carbon-based molecules were excluded from the stoichiometric matrix, such as H2O, ATP, NADH, or other co-factors (see complete list in Supplementary Table S1). We further reduced the complexity of the factor graph based on its topological structure. In this step, connected reactions were merged into a module if (1) none of the merged intermediate metabolites has more than one in-flux or out-flux reactions that correspond to more than one module inputs or outputs; and (2) none of the merged intermediate metabolites has an in-flux or out-flux other than merged reactions or the module input and output. We have proved that under these two conditions and the flux balance condition, changes of the reactions inside the module will not affect the reactions outside of the module conditional on a fixed flux rate of the module, i.e., solving the flux of each individual reaction in a merged module is equivalent to solve the flux of the module (see details in Supplementary Methods). Noted, the merged reactions will form a variable node containing multiple reactions in the factor graph, while the factor nodes are still individual metabolites. In addition, we identified certain classes of metabolites, including different types of fatty acids, pyrimidines, purines, and steroid hormones, form highly connected web-like metabolic pathways. Instead of solving the flux for each individual metabolite, we consider the metabolites of the same class as one factor. The network reduction approach enables a more robust flux estimation by estimating the flux of one module instead of individual reactions, and a more efficient computation over the simplified network topological structure.

We reconstructed the human metabolic map into a factor graph consisting of 169 modules of 22 super module classes, 862 genes, and 128 metabolites, out of which 66 are intermediate substrates. Here each super module is a manually curated group of modules of a similar function (**Table 1**). Detailed information of the factor graphs is listed in **Table 1** and Supplementary Table S1. **Figure 2A** illustrates the functional group and complete topological structure of the collected metabolic modules and super modules in human. **Figure 2B** shows the reconstructed factor graph for human metabolic map, which is utilized in further flux estimation. **Figure 2C** illustrates several examples of how network motifs in the input metabolic network are merged into one metabolic module.

**Figure 2.**
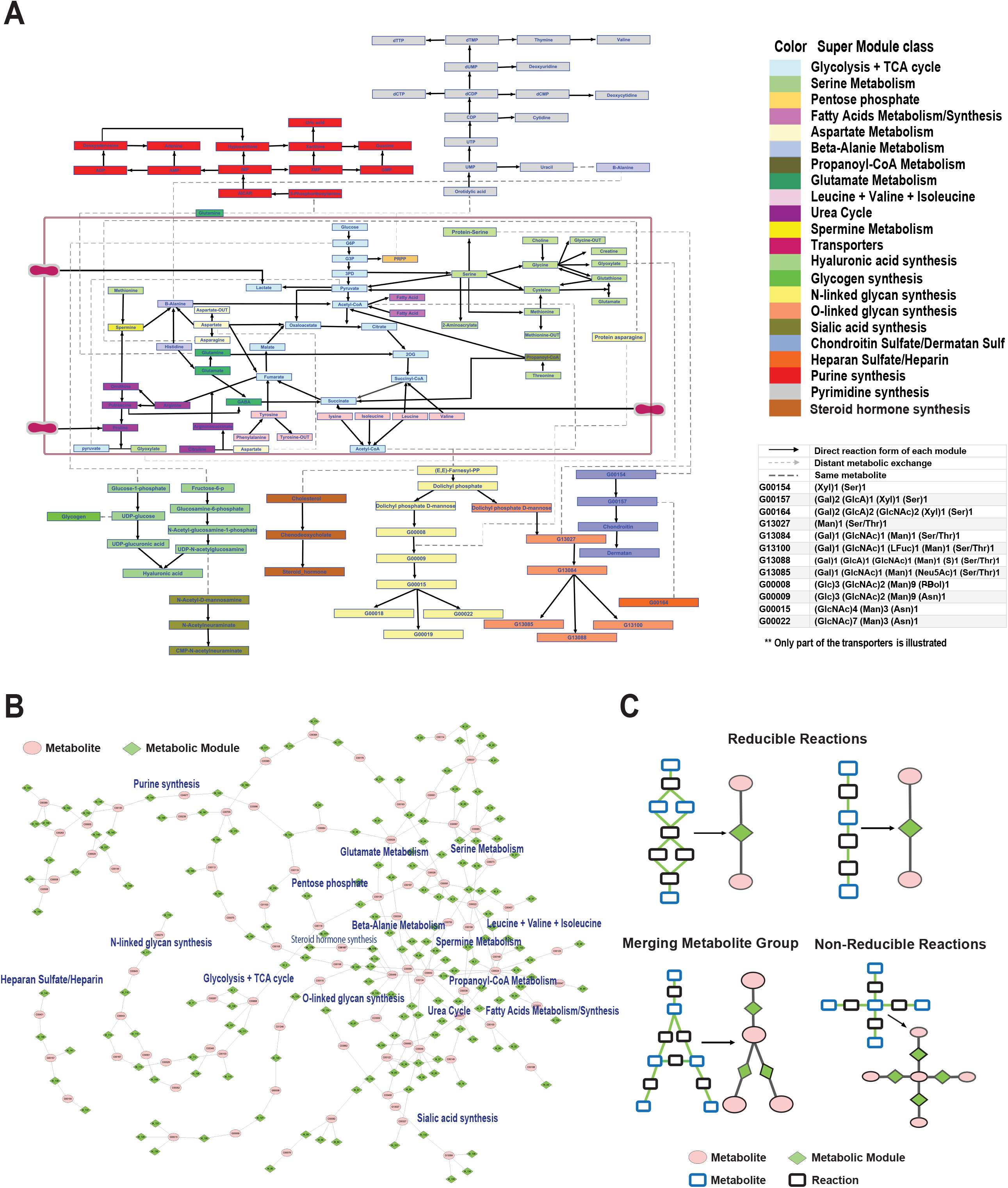
Reduced and reconstructed human metabolic map. (A) Collected human metabolic modules and super module classes. (B) Factor graph representation of the reconstructed human metabolic map, in which the modules and metabolites were colored by green and pink, respectively. (C) Examples of how the network motifs in the metabolic map are simplified into metabolic modules, where the reactions and metabolites are represented by black and blue rectangular, and modules and metabolites are colored by green and pink. Chain-like reactions can be directly simplified; a complicate module connected by multiple branches can be shrunk into one point linked with the multiple branches; and complicated intersections cannot be simplified.

For a given scRNA-seq data and a user defined task, scFEA further refine the task specific metabolic factor graph by: (1) limiting the analysis to user selected metabolic networks, and (2) exclude the modules without significantly expressed gene. For (2), scFEA will first determine for all the genes whether they have a significant non-zero expression state using our in-house Left Truncated Mixture Gaussian model (Wan et al. 2019b) (see details in Methods). Under the default setting of scFEA, a module is considered as blocked if it becomes disconnected after removing the reactions whose associated genes do not have a non-zero expression state. The blocked modules will be excluded from further analysis. On account of the common drop-out events in scRNA-Seq data, scFEA also enables a more conservative assumption to remove a module only if none of the genes involved in all reactions of this module has significantly active expressions.

The topological structure of metabolic modules including input, output and intermediate metabolites and genes associated with each module will be utilized for further flux estimation.

### Mathematical formulation of metabolic flux estimation in individual cells

For a clear model setup, we first formulate the metabolic network as a directed factor graph, in which each metabolic module is represented as a variable, each compound is represented as a factor node carrying a loss function to evaluate the level of flux balance among modules, and the direction represents if a metabolite is the input or output of a metabolic module (**Figure 2B**). We denote *FG*(*C*^*1×K*^,*RM*^*1×M*^,*E* = {*E*_*C→R*,_*E*_*R*→C_}) as the factor graph, where *C*^1×K^ = {C_K,_ *k* = 1,…*K*} is the set of *K* compounds, *RM*^1×*M*^ = {*R*_*m*,_*m* = 1,…,*M*} is the set of *M* metabolic modules *E*_*C→R*_ and *E*_*R→C*_ represent direct edges from module *R*_*m*_ to compound *C*_*K*_ and from compound *C*_*K*_ to module *R*_*m*_ respectively. For the *k*-th compound *C*_*k*_, we define the set of reactions consuming and producing *C*_*k*_ as 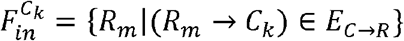 and 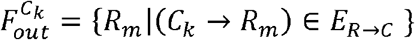, which is derived from the stoichiometric matrix of the whole metabolic map. For an scRNA-seq data set with *N* cells, we denote *Flux*_*m,j*_ as the flux of the *mth* module in the cell *j, j* = 1…*N* and *F*_*j*_ *=* {*Flux* _1,j, …,_ *Flux* _*M,j*_} as the whole set of the reaction fluxes. Our computational hypothesis is that total flux imbalance of the intermediate metabolites though all the collected cells should be minimized, based on which we developed the likelihood function of the flux of all modules through all cells as:

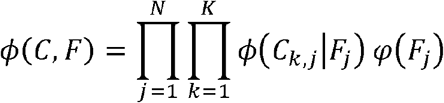

where 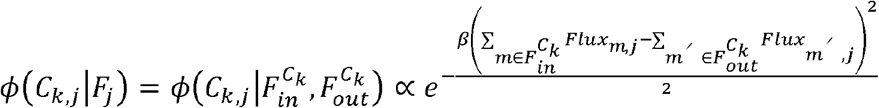 and φ(*F*_*j*_) represents the prior distribution of the fluxome in cell *j*, and *β* is a tuning hyperparameter. scFEA models the flux of reach reaction,*Flux*_*m,j*,_ as a nonlinear function of the expression changes of the genes associated with the module. Denote 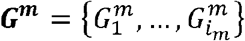 as the genes associated with the reactions in *R*_*m*_, and 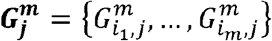 as their expressions in sample *j*, where *i*_*m*_ stands for the number of genes in *R*_*m*_. We model 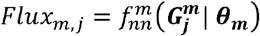 multi-layer fully connected neural network with the input 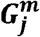, where ***θ***_***m***_ denotes the parameters of the neural introduced network (**Figure 3**). It is noteworthy that the cell group and tissue context specific distribution of the flux *φ*(*F*_*j*_) and the reaction parameters ***θ***_***m***_, are always unknown. Here the flux balance assumption leads to a self-constrained loss function and *Flux*_*m,j*,_ ≡ 0 forms a general trivial solution. To provide a robust and rational solution, We indroduced two additional assumptions to *Flux*_*m,j*,_, namely (1) the predicted flux should be non-negative and (2) the total predicted flux in each large super module tends to have a high correlation with gene expression variation of the super module (**Figure 2A**). The second assumption considers the metabolic flux variation of large metabolic modules among single cells is coherent to their gene expression change, which could be supported by several recent studies (Damiani et al. 2019a; Wagner e t al. 2 020). In addition, this assumption effectively avoids the trivial solution. Hence, we instead of directly maximize *ϕ*(*C,F*), we solve the ***θ***_***m***_ and cell-wise flux *Flux*_*m,j*_ by minimizing the following *L*:

**Figure 3.**
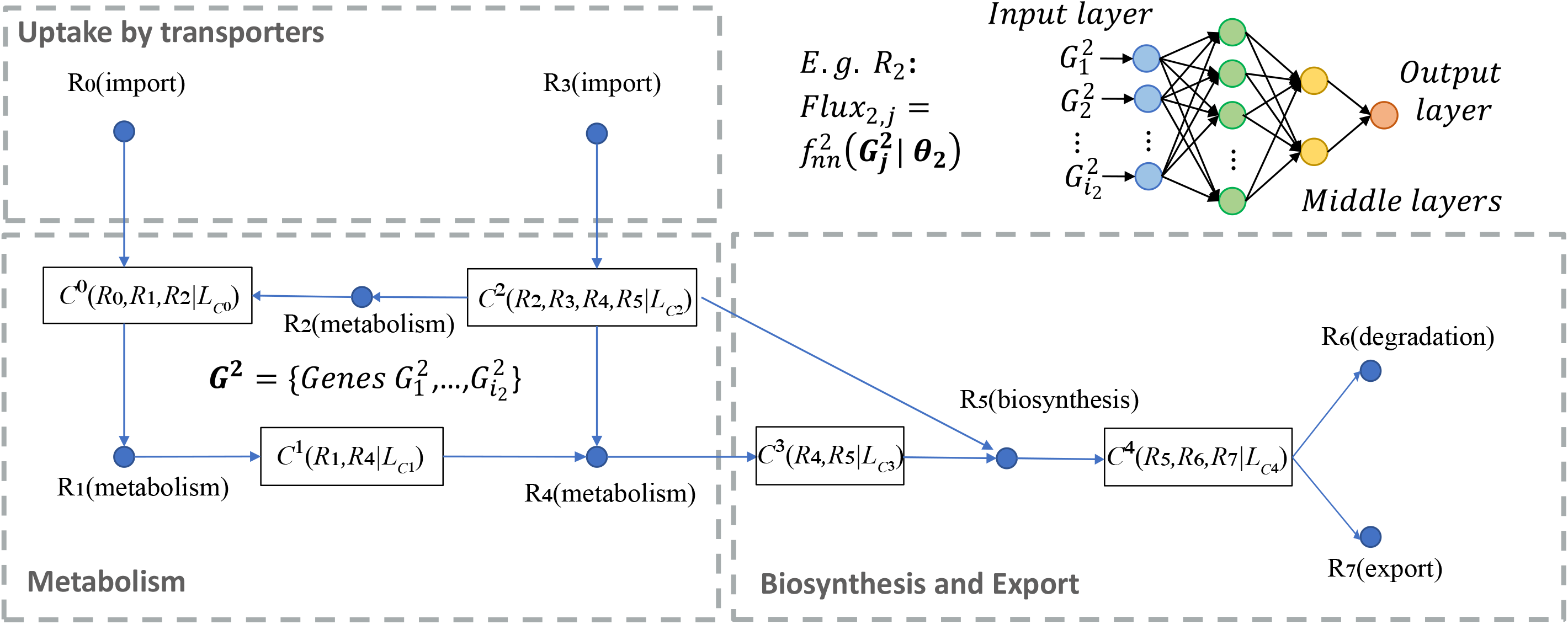
A toy model of the factor graph of metabolic modules, flux balance conditions, and the flux model for the module *R*_2_ (top-right). In the factor graph, each C (metabolites) corresponds to one flux balance condition and serves as a factor, and each R (can be a reaction or a module) is a variable. For example 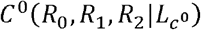 simply represents that the metabolite *C*^0^ is determined by the flux balance loss of *R*_0_ *R*_1_ *R*_2_, here 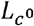 is the flux balance term of *C*^0^. Import and export/degradation reactions are considered as having no input or output substrates.

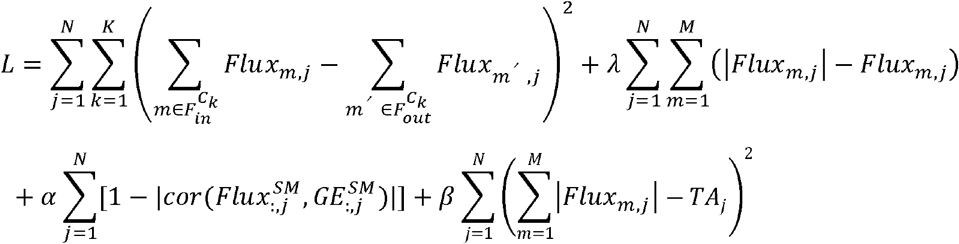

where *λ, γ, α* and *β* are hyperparameters, *cor* represents Pearson correlation coefficients; *Flux*^*SM*^ and *GE*^*SM*^ are two *NSM* x *N* two matrices, here *NSM* is number of super modules, 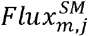 represents the sum of the flux of the modules in the super modules *m*, 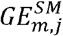 represents the sum expression of the genes in the super module *m*, in cell *j*, and *TA*_*j*_ is a surrogate for total metabolic activity level of cell *j*, which is assigned as the total expression of metabolic genes in cell *j*. The first, second, third and fourth term of *L* are the loss of flux balance, non-negative flux, the coherence between predicted flux and total gene expression level of each super-module, and flux scale. Noting genes may have varied intrinsic expression range, decentralized Pearson correlation is utilized to model the coherence between gene expression and predicted flux.

It is noteworthy that the above formulation defines a new graph neural network architecture for flux estimation over a factor graph, where each variable is defined as a neural network of biologically meaningful attributes, i.e., the genes participating in each metabolic module, and the information aggregation between adjacent variables is constrained by the balance of chemical mass of the in- and out-flux of each intermediate metabolites. Noted, the number of intermediate constraints (*K*) and large sample size (*N*) of scRNA-seq data ensures the identifiability of ***θ***_***m***_ for the 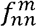 at a certain complexity level. Another advantage of this formulation is that the flux balance loss forms a self-constraint term that takes chemical stoichiometric into consideration. In addition, our formulation does not require a prior knowledge of the imports and exports of the whole system, which are always cell and context specific and unknown. Detailed analysis of the computational feasibility, scalability, tuning of hyperparameters, and options of the additional loss terms are provided in Methods and Discussions.

The challenges to minimize the loss function *L* include the following: (1) the flux of each module affects the balance of its input and output and multiple modules are involved in the balance of one intermediate substrate, hence perturbing one single flux at each step may not converge, and on the other hand (2) the direction for simultaneously updating a large group of fluxes cannot be theoretically derived. The two challenges prohibit a direct utilization of back propagation or gradient descending methods. We developed an effective optimization strategy for by adopting the idea of information transfer in belief propagation, which has been commonly utilized in analyzing cyclic networks such as Markov random field (Lan et al. 2006). Specifically, *L* is minimized by iteratively minimizing the flux balance of each intermediate metabolite *C*_*K*_ and the weighted sum of the flux balance of the Hop-2 neighbors of *C*_*K*_ in the factor graph, as the 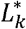 defined below:

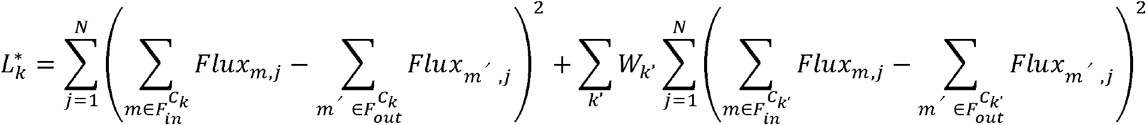

where C_*k*′_are the Hop-2 neighbors of C_*k*′_, *W*_*k*′_ is proportional to the current total imbalance of all the Hop-2 neighbors of *C*_*k*′_ except for *C*_*k*_ itself (see more details in Methods). Here the Hop-2 neighbors of a compound (or module) on the factor graph is defined by all other compounds (or modules) having a connection with the modules (or compounds) who connect to the compound (or module). Such a regional perturbation strategy over the whole graph can effectively leverage the search of global minimum and computational feasibility.

The output of scFEA includes 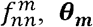, ***θ***_***m***_ for each module and predicted cell-wise metabolic flux *Flux*_*m,j*._ noteworthy the predicted flux *Flux*_*m,j*_ is a relative measure of unfixed scale. However, *Flux*_*m,j*_ is comparable among cells (*Flux*_*m*,_) or metabolic modules (*Flux*_,*j*_).

### Method validation on a scRNA-seq data with perturbed metabolic conditions and matched metabolomics data

To validate the cell-wise flux estimated by scFEA, we generated an scRNA-seq dataset consisting of 162 patient-derived pancreatic cancer cells (Pa03c cell) under two crossed experimental conditions: APEX1 knockdown (APEX1 KD) or control, and under hypoxia or normoxia conditions (see detailed experimental procedure and data processing in Methods). Metabolomics profiling of 14 metabolites, namely glucose, glucose-1 phosphate, glucose-6 phosphate, pyruvate, and lactate in the glycolysis pathway, citrate, 2-oxoglutarate, succinate, fumarate, malate in the TCA cycle, and amino acids glutamate, glutamine, serine, and ornithine were collected on bulk wildtype Pa03c cells and APEX1 inhibition cells under the normoxia conditions, each with three replicates (Supplementary Table S2). We utilized the Smart-seq2-fluidigm protocol for single cell RNA sequencing for saturated gene detection of each single cell, to enable a more accurate modeling of metabolic flux. *APEX1* is a multifunctional protein that interacts with multiple transcriptional factors (TFs) to regulate cellular responses to hypoxia and oxidative stress (Kelley et al. 2012). Our previous studies identified significant roles of *APEX1* in the regulation of Pa03c cells’ response to metabolic environment changes (Shah et al. 2017; Wan et al. 2019a).

To the best of our knowledge, scFEA is the first computational tool to estimate metabolic flux at single cell level. Without baseline methods for comparisons, we validate scFEA by examining the consistency between the metabolic flux variation predicted by scFEA and experimental observations. We identified 126 up- and 443 down-regulated genes in APEX1 KD vs Control under the normoxia condition, and 260 up- and 1496 down-regulated genes under hypoxia condition. Pathway enrichment analysis showed that the TCA cycle pathway (normoxia: *p*=0.003, hypoxia: *p*=1.12e-07) and oxidative phosphorylation (normoxia: *p*=3.17e-4, hypoxia: *p*=1.77e-08,) are significantly enriched by down regulated genes, under both normoxia and hypoxia conditions. This suggests that the knock down of APEX1 may lead to inhibited cellular aerobic respiration. In addition, genes regulated by *HIF1A* (hypoxia-inducible factor 1-alpha), including glycolysis and TCA cycle genes, were observed to be up- and down-regulated respectively, in comparison to the hypoxia vs normoxia conditions in the control Pa03c cells. This is consistent to the common knowledge of hypoxia response. Out of the 14 metabolites, we have seen increase of abundance in glucose, glucose-1 phosphate, glucose-6 phosphate, and lactate, and decrease in 2-oxoglutarate, succinate, fumarate, and malate in APEX1-KD vs control cells under the normoxia condition. In summary, analysis of the single cell gene expression and bulk cell metabolomic data revealed that knockdown of APEX1 affects the cells’ glucose metabolism and inhibits the cells’ TCA cycle pathway, under both normoxia and hypoxia condition. **Figure 4A** illustrates the variation of genes and metabolites involved in glycolysis, pentose phosphorylation, TCA cycle, glutaminolysis and aspartate metabolism pathways in APEX1-KD vs control under normoxia condition. We conducted a qRT-PCR experiment to confirm the down regulated genes in glycolysis, TCA cycle and oxidative phosphorylation pathways (Supplementary Figure S1). Complete list of differentially expressed genes and pathway enrichment results were provided in Supplementary Table S3.

**Figure 4.**
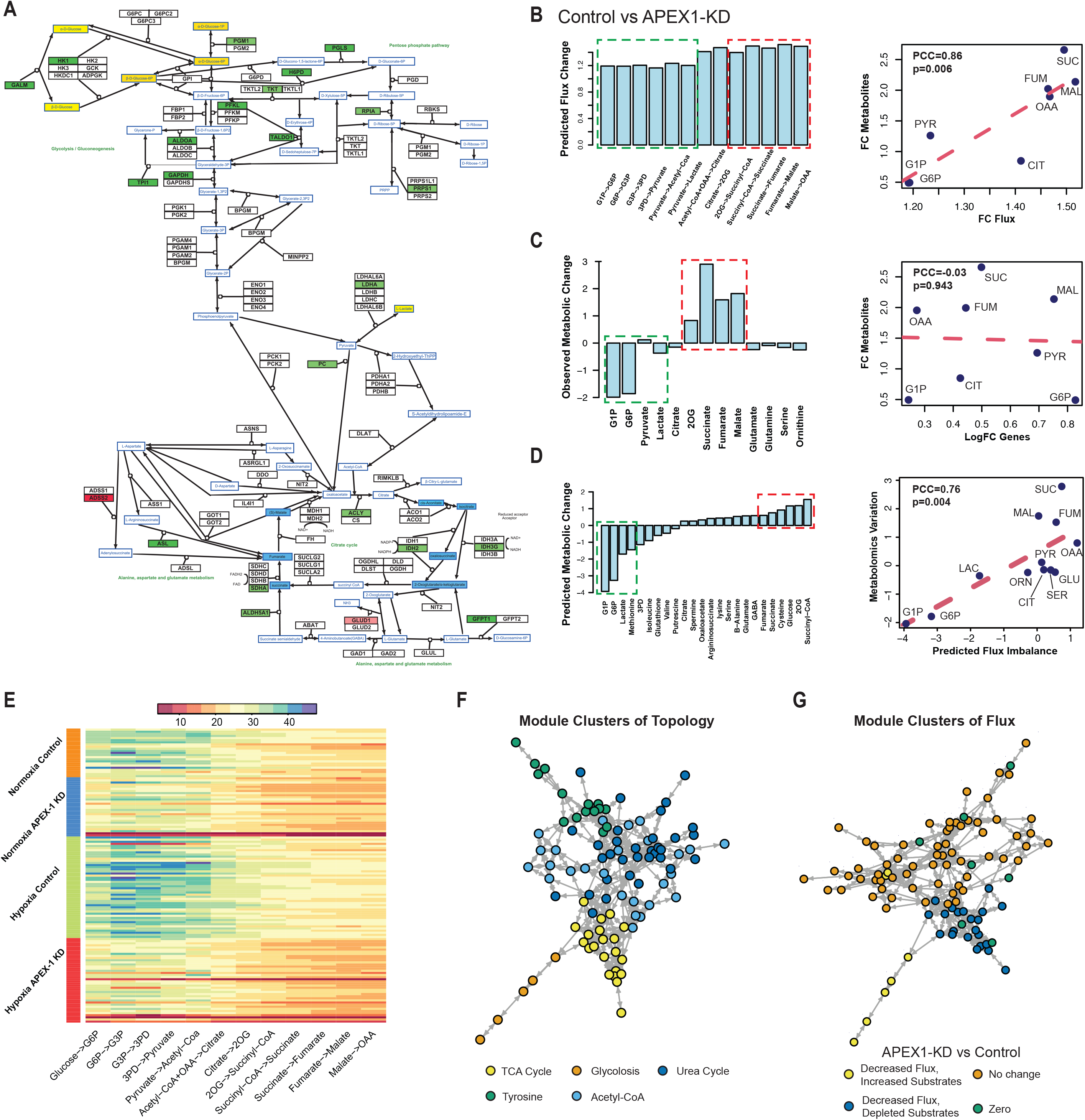
Application of scFEA on matched scRNA-seq and metabolomics data of Pa03C cells. (A) Gene expression and metabolomic variations of the glycolysis, pentose phosphate, TCA cycle, glutamine, and aspartate metabolic pathways in APEX1-KD vs control under normoxia condition. Genes/metabolites were shown in rectangular boxes with black/blue borders, up/down regulated genes were colored in red/green, increased and decreased metabolites were colored in yellow/blue, respectively. The darker color suggests a higher variation. (B) Predicted flux fold change (left, *x*-axis: metabolic module, *y*-axis: predicted flux change) in control vs APEX1-KD, and correlation between fold change of predicted flux and observed metabolite change (right, *x*-axis: fold change of predicted flux, *y*-axis: fold change of observed metabolite abundance, each data point is one metabolite, PYR: pyruvate, CIT: citrate, FUM:fumarate, SUC: succinate, MAL: malate). (C) Observed metabolomic change (left, *x*-axis: metabolites, y-axis: abundance difference observed in the metabolomics data) in control vs APEX1-KD, and correlation between log fold change of gene expressions involved in each reaction and observed metabolomics change (right, *x*-axis: log fold change of the averaged expression of the genes involved in each reaction, *y*-axis: fold change of observed metabolites abundance observed in the metabolomics data, each data point is one metabolite). (D) Predicted metabolic stress (left, *x*-axis: metabolites, y-axis: predicted abundance difference) in control vs APEX1-KD and correlation between predicted metabolic stress and observed difference in metabolite abundance (right, *x*-axis: top scFEA predicted imbalance of the in-/out-flux of intermediate metabolites, *y*-axis: difference of observed metabolomic abundance, in control vs APEX1-KD, each data point is one metabolite: LAC: lactate, SER: serine, GLU: glutamine, ORN: ornithine). In (B-D) all comparisons were made by comparing control vs APEX1-KD under normoxia. Noted, the fold change of metabolomic abundance is used in calculating the correlation in B-C and difference of metabolomic abundance is used in D. The green and red dash-blocks represents the accumulated (green) and depleted (red) metabolites in Control vs APEX1-KD. (E) Profile of the predicted fluxome of 13 glycolytic and TCA cycle modules. Here each column represents the flux between two metabolites, shown on the *x*-axis, for all the cells of the four experimental conditions, shown on the *y*-axis. For two neighboring fluxes, the product of the reaction on the left is the substrate of the reaction on the right, and in a perfectly balanced flux condition, the two neighboring fluxes should be equal. (F) Clusters of metabolic modules inferred by using the network connectivity structure only. (G) Clusters of metabolic modules inferred by using the network topological structure (weight of 0.3) combined with predicted fluxome (weight of 0.7).

#### Consistency between the scFEA predicted flux variation and the metabolomics data

We applied scFEA to the aforementioned scRNA-seq data of the four conditions. We first focus on the normoxia conditions where matched single cell expression and metabolomics data are available. scFEA predicted decreased metabolic flux for the modules in glycolysis and TCA cycle in APEX1-KD vs control, i.e., glucose → D-Glucose 1-phosphate (G1P) → alpha-D-Glucose 6-phosphate (G6P) → glyceraldhyde-3P (G3P) → 3-Phospho-D-glyceroyl phosphate (3PD) → pyruvate → Acetyl-CoA → citrate → 2-Oxoglutarate (2OG) → succinate-CoA → succinate → fumarate → Malate → oxaloacetate (OAA) and pyruvate lactate, where particularly, the reactions towards the downstream of this reaction chain has even lower flux in APEX1-KD (**Figure 4B**). We examined the correlation between the averaged predicted flux change with the observed metabolomic change of intermediate metabolites in glycolysis and TCA cycle pathways in APEX1-KD vs control cells under normoxia condition and observed a Pearson Correlation Coefficient (PCC) of 0.86 (*p*=0.006) (**Figure 4B**), suggesting the high consistency between predicted flux variation with the observed metabolic changes. Using metabolomics data, we observed increase of production for glucose, G1P, G3P and lactate, and decrease of production for 2OG, succinate, fumarate, and malate in APEX1-KD vs control (**Figure 4C**). By Michaelis Menten model, the substrates of largely varied concentration determine the reaction rate in a non-linear manner (close to linear when the reaction is less saturated). Hence, variations in the concentration of the metabolites with one dominating out-flux could partially reflect the changes of the out-flux rate. We also correlated the metabolomic change with the averaged expression change of the enzymes catalyzing the reactions. However, no significant correlation was observed (PCC=-0.03, p=0.943, **Figure 4C**), suggesting that single cell gene expression itself, without considering the constraints from the intricate metabolic network as in scFEA, doesn’t produce a good estimate of single cell metabolic landscape. In addition, ssGSEA (single sample gene set enrichment analysis) has been utilized to model cell-wise pathway activity in scRNA-seq data (Chen et al. 2020). The correlation between the metabolomic changes and the differences in averaged ssGSEA score in APEX1-KD vs control cells is (PCC=0.42, p=0.299) (Supplementary Figure S2). Here scFEA leveraged the non-linear relationships between gene expression and enzymatic reaction rate, and the flux balance constraints of the metabolites, hence its predicted metabolic flux is more consistent to the true metabolomics changes.

#### High consistency of the predicted metabolic stress with experimentally observed metabolomic changes

scFEA predicted in and out flux for each metabolite allows us to investigate the cell-wise metabolic stress, which was defined as the imbalance of the in-/out-fluxes of each intermediate metabolites in each cell. **Figure 4D** shows that the G1P, G6P and lactate were accumulating while 2OG, succinate, succinyl-CoA, and fumarate were depleted in APEX1-KD vs control. A PCC of 0.75 (*p*=0.004) was observed between the predicted metabolic stress and the true metabolic change, on 12 metabolites with both measured metabolomic profile and predicted metabolic stress, demonstrating a high accuracy of the predicted metabolic stress level. Detailed predicted and observed metabolic imbalance were provided in Supplementary Table S2. **Figure 4E** shows the predicted cell-wise fluxome of the glycolysis and TCA cycle modules for cells of the four conditions. We observed, in general, higher flux of the glycolytic modules than the TCA cycle modules, with the largest average flux gap seen on Pyruvate → Acetyl-CoA and Acetyl-CoA → Citrate.

In addition, the flux of the downstream reactions (citrate → 2OG → succinyl-CoA → succinate) of the TCA cycle is lower than the upstream reactions (succinate → fumarate → malate → OAA). A possible explanation for the leaky metabolic flux is that some of the intermediate substrates’ flow to other branches, majorly for biosynthesis of amino acids. Among the four conditions, we identified that the hypoxia control group has the highest flux rate of glycolysis and TCA cycle modules. Clearly, the inhibition of APEX1 significantly decreased the metabolism rate of glucose. Combined with the accumulations of glycolytic substrates and depletions of TCA cycle substrates identified by the metabolic stress and metabolomics data analysis, our speculate that the knock-down of APEX1 may directly impact the downstream part of glycolysis, the whole TCA cycle and further oxidative phosphorylation, leading to accumulation of G1P and G6P as a result of the blockage. Up regulation of glucose transporters was also observed in APEX1 KD vs control, further suggesting the accumulation of glycolytic substrates.

#### Perturbation analysis of flux deterministic genes

We also conducted a perturbation analysis to tease out the key genes with high impact on each metabolic module (see details in Methods). The following genes were identified to have the highest impact on metabolic flux: HK1 and HK2 (Glucose→G6P, EC: 2.7.1.1), ALDOA, PFKL and GPI (G6P→G3P, EC: 5.3.1.9), GAPDH and PGK1 (G3P→3PD, EC: 1.2.1.12, 2.7.2.3), ENO1, PGAM1, and PKM (3PD→Pyruvate, EC: 5.4.2.11, 4.2.1.11), PDHA2 (Pyruvate→Acetyl-Coa, EC: 1.2.4.1), LDHA (Pyruvate→Lactate, EC: 1.1.1.27), ACLY (Acetyl-CoA+OAA → Citrate, EC: 2.3.3.8), IDH2 (Citrate → 2OG, EC: 1.1.1.42), DLD and OGDH (2OG→Succinyl-CoA, EC: 1.2.4.2), SUCLG1 (Succinyl-CoA→Succinate, EC: 6.2.1.4), SDHA (Succinate→Fumarate, EC: 1.3.5.1), FH (Fumarate→Malate, EC: 4.2.1.2), MDH1 (Malate→OAA, EC: 1.1.1.37). Detailed results of the perturbation analysis were illustrated in Supplementary Figure S3. A qRT-PCR experiment was conducted to confirm the down regulation of the above key metabolic genes, including HK1, PFKL, ACLY, SDHA, and IDH2 (Supplementary Figure S1). We also correlated the predicted high impact enzyme in the modules containing multiple enzymes (seven in total) with the rate limiting enzymes reported in Rate-Limiting Enzymes database (RLEdb) (Zhao et al. 2009). We observed the six out of the seven predicted high impact enzymes, namely 2.7.1.1, 1.2.1.12, 2.7.2.3, 5.4.2.11, 1.2.4.1, and 1.2.4.2, have been reported, suggesting a significant enrichment (p=0.0005 by Fisher’s exact test) of our predictions to previously reported rate limiting enzymes. We further conducted a module level perturbation analysis by increasing or decreasing the expression of genes in a certain module (Methods). Non-surprisingly, a decrease of expression on genes of the downstream part of glycolysis pathway in the control cells will lower the flux of the TCA cycle, causing the accumulation of glycolytic intermediate substrates and depletion of TCA cycle metabolites, which is consistent to our experimental observations.

#### Detecting groups of metabolic modules with similar variations and cells with distinct metabolic states

We also applied scFEA to a larger metabolic map, with the 11 metabolic super modules and transporters, and then examined the high-level organization of the modules. Based on only the metabolic network connectivity, **Figure 4F** illustrated five distinct groups of metabolic modules derived using a spectral clustering method purely based on their network topology (see Methods), namely (1) glycolysis, (2) TCA cycle and glutamine metabolism related modules, (3) tyrosine and serine metabolism, (4) urea cycle related modules, and (5) acetyl-coA related metabolisms such as fatty acids and propanoyl-CoA metabolisms. To examine the high level structure based on the flow of flux, we further conducted a clustering analysis of the metabolic modules by considering both the network connectivity and flux similarity. The distance between two modules *R*_*i*_ and *R*_*j*_ is defined as *αd*(*R*_*i*_, *R*_*j*_) + (1 − *α*)*d*^*F*^(*R*_*i*,_ *R*_*j*_), where *d*(*R*_*i*,_ *R*_*j*_) is their normalized spectral distance, and *d*^*F*^(*R*^*i*,^ *R*^*j*^), is their normalized similarity in estimated flux through all the normoxia cells (see Methods). Here α = 0.3 is used in the analysis. **Figure 4G** shows the metabolic module clusters by integrating topological structure and flux similarity. Four distinct clusters were identified, including (1) glycolysis and fatty acids metabolism of decreased flux and accumulated substrates in APEX1-KD vs control, (2) TCA cycle and pyruvate metabolism decreased flux and depleted substrates, and (3) metabolism of amino acids and other metabolites with unchanged flux and metabolites, and (4) a few other modules of 0 flux rates, respectively. This observation further validated the rationale of scFEA predicted fluxome.

We also conducted cell clustering based on the metabolomic modules with varied flux (Methods). Non-surprisingly, the cells clusters were aligned with experimental conditions, forming five group of cells of high, intermediate, and low, low metabolic rates, high lactate production and low TCA-cycle (Supplementary Figure S4).

### Method validation and robustness analysis on synthetic and independent real-world data

#### Method validation on independent real-world data

We also validated scFEA on an independent scRNA-seq data of perivascular adipose tissue derived mesenchymal stem cells (PV-ADSC) (GSE132581) (Gu et al. 2019). To the best of our knowledge, this data set plus our newly generated data sets are the only two scRNA-seq data with matched tissue level targeted metabolomics profiling available in the public domain. We first re-conducted cell clustering analysis and identified two distinct cell clusters corresponding to the PV-ADSC of different level of differentiation, as reported in the original work (**Figure 5A**). Due to the small sample size (85 cells), scFEA was applied to estimate the fluxome of glycolysis and TCA cycle pathways. We observed an increased flux of glycolytic reactions (p<1.56e-6), lactate production (p=0.002), and the reactions from cis-aconitate to oxaloacetate in TCA cycle (p<0.02) in the more differentiated (MD) PV-ADSC vs the PV-ADSC of high stemness (HS) while the reactions from acetyl-CoA to citrate were insignificantly changed (p=0.887) (**Figure 5B**). Our predicted flux variations between the two cell clusters are consistent to the metabolomic observations made in the original work, i.e., the glycolytic intermediate metabolites, lactate production, and metabolites in the later part of TCA cycle were evaluated in the more differentiated PV-ADSC and citrate was not significantly changed. We also analyzed the metabolic modules of two amino acids super modules with metabolomics profile reported in the original study, namely valine and isoleucine metabolism and glutamate and glutathione metabolism (Supplementary Figure S5). Elevated valine and isoleucine metabolic flux in MD vs HS PV-ADSC has been predicted by scFEA, which is consistent to the original report. scFEA also predicted an increased flux of the modules from glutathione → glutamate → glutamine → TCA cycle, this could explain the increased flux rate of TCA cycle but less increase in citrate production. The original study only reported a depletion of glutathione and glutamate, our metabolic stress analysis also predicted more decreased glutathione and glutamate in MD vs HS PV-ADSC. Our analysis suggested the evaluated glutamate and glutathione metabolism is to fuel the substrate source for TCA cycle in MD cells, which depleted the concentration of glutathione and glutamate.

**Figure 5.**
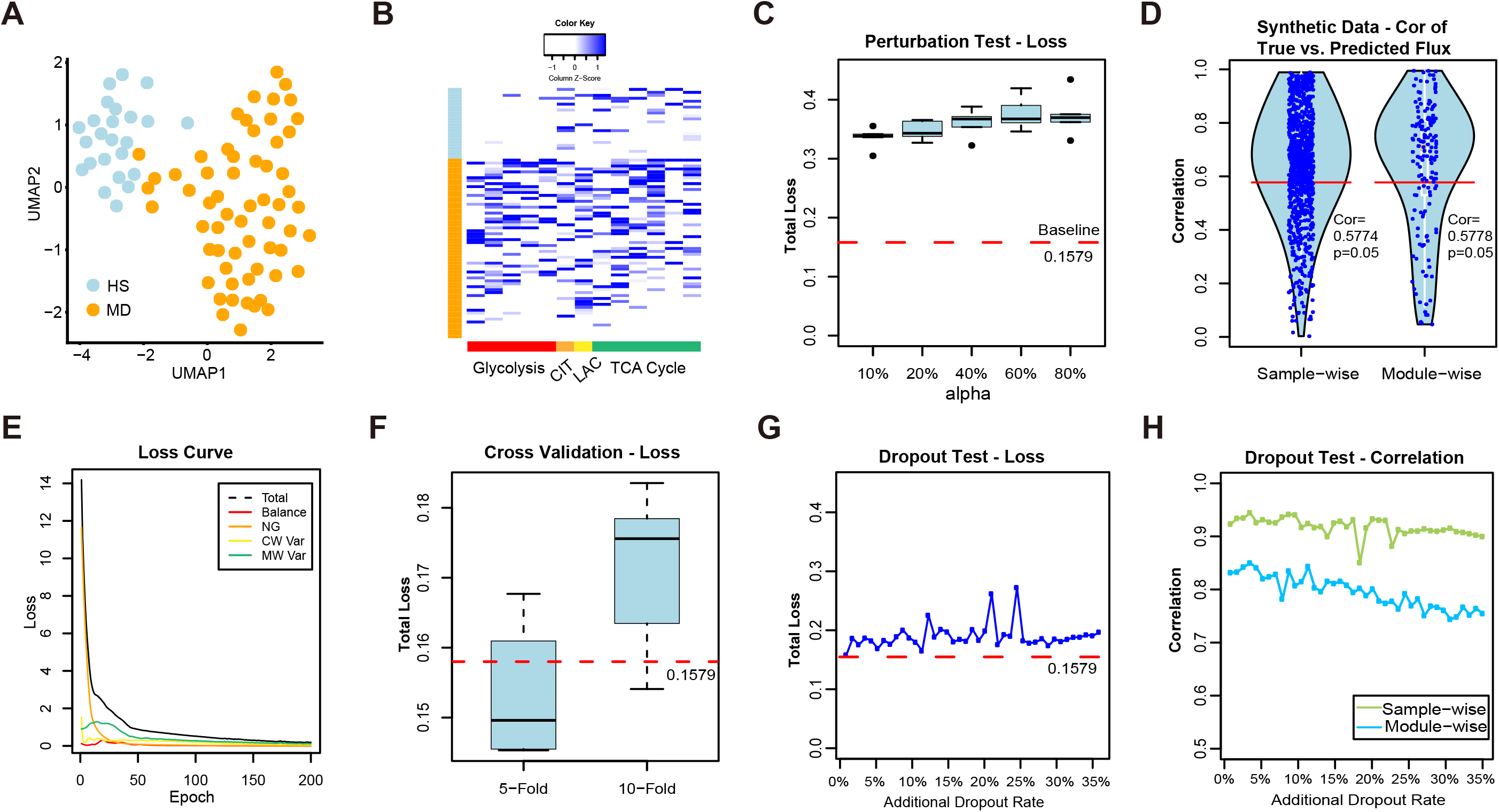
Methods validations on real-world and synthetic datasets. (A) UMAP of the cell clustering analysis of the GSE132581 PV-ADSC data, here HS and MD stand for PV-ADSC of high stemness and more differentiation, respectively. (B) Distribution of predicted cell-wise flux of glycolytic and TCA cycle modules. Each row is one cell, where row side color bar represents HS and MD PV-ADSC by blue and orange, respectively. Each column is one module. The left five columns (red labeled) are glycolytic modules from glucose to acetyl-CoA, the CIT column (orange labeled) is the reaction from acetyl-CoA to Citrate, the LAC column (yellow labeled) is the reaction from pyruvate to lactate, and the right six columns (green labeled) are TCA cycle modules from citrate to oxaloacetic acid. (C) Total loss (y-axis) with respect to different proportion of cell samples (x-axis) with randomly shuffled gene expressions of the pancreatic cancer cell line data. The baseline loss 0.1579 was computed by using the original expression profile of all 166 cells. (D) The sample-wise and module-wise correlation (y-axis) between the true and predicted module flux in synthetic data-based method validation, here Cor=0.5775 (p=0.05) and 0.5778 (p=0.05) correspond to the sample-wise and module-wise correlation, respectively. (E) Convergency of the total loss and four loss terms during the training of neural networks on the pancreatic cancer cell line data. (F) Total loss (y-axis) computed under 5-/10-fold cross validation (x-axis) vs baseline loss. (G) Total loss (y-axis) computed from the robustness test by adding 0%-35 artificial dropouts to the original data (50.22% zero rate) vs baseline loss. (H) Sample-wise and module-wise correlation (y-axis) of the module flux predicted from the data with 0%-35 additional artificial dropouts with the module flux predicted from the original data.

#### Method validation on randomly shuffled gene expression profile

In scFEA, we assume that the flux distribution in each single cell should be constrained by the flux balance condition while the reaction rate of each module could be modeled as a non-linear function of the gene expressions involved in this module. These two assumptions suggested that the distribution of the gene expressions involved in the metabolic modules was essentially determined by the metabolic flux distribution and constrained by the flux balance condition. One existing evidence directly supports our assumptions is that the expression of closely related metabolic genes always tend to be co-up or co-down regulated (van der Knaap and Verrijzer 2016; Li et al. 2018). To further validate our assumption, we randomly shuffle the expression profile of each gene in a certain proportion (10%, 20%, 40%, 60% and 80%) of cells in our pancreatic cancer cell line data, and applied scFEA to each perturbed data (see details in Supplementary Methods). We observed that the minimized total loss is positively associated with the level of perturbations (**Figure 5C**) and the original scRNA-seq data achieved the smallest total loss, which partially support our underlying assumption.

#### Method validation on synthetic data

We also conducted a synthetic data based experiment and demonstrated that the loss function and solution strategy of scFEA can accurately estimate cell-wise metabolic flux from scRNA-seq data. We first randomly generated 1000 cells having different flux distribution of 169 connected modules from the solution space satisfying flux balance condition of these modules. The expression profile of the genes involved in each module was reversely simulated by assuming its flux follows a fixed non-linear function of the gene expressions. Two levels of errors were added to the flux and gene expression level. Detailed data simulation approach was provided in Supplementary Methods. We applied scFEA on the simulated single cell gene expression profile and compared the fluxome predicted by scFEA and known fluxome. We observed that scFEA predicted fluxes are highly consistent to the true flux distribution, on both directions of the cells and metabolic modules (**Figure 5D**). Specifically, more than 70% single cells achieved at least 0.5774 (p=0.05) sample-wise correlation and more than 70% modules achieved at least 0.5778 (p=0.05) module-wise correlation. Our analysis demonstrated that under the assumption of scFEA, i.e., if the flux balance constraint and non-linear dependency between gene expression and metabolic hold, the formulation and solution strategy of scFEA could accurately estimate the cell-wise fluxome from single cell gene expression data.

#### Robustness analysis based on perturbed sample inputs, cross-validation, and analysis of hyperparameters

We also tested the robustness of scFEA by 2-/5-/10-fold cross validations on the pancreatic cancer cell line data. Our analysis suggested that both total loss of the testing data does not change significantly when the number of input cells varied from 50% to 100% of all the cells (**Figure 5E**). For hyperparameters tuning, scFEA used Adam as the optimizer, which can automatically adjust the learning rate. To choose the most suitable hyperparameters of the four terms in the loss function, we conducted experiments by changing the relative scale of any two terms and fixing the rest two on the pancreatic cancer cell line data. We update two hyperparameters relative ratio from 10 to 1000. Our experiments suggested a similar optimal solution can always be achieved under our hyperparameter perturbation range (Supplementary Figure S6). **Figure 5F** showcases the convergency of the four loss terms and total loss in the model fitting of the pancreatic cancer cell line data. In addition, the applications on six real-world data (see further results) and simulated data suggested that the default hyperparameters always generate results of good convergency of the total loss and high biological implications. The default hyperparameters of the current version and details in hyperparameter tunning codes were provided via https://github.com/changwn/scFEA.

#### Robustness analysis with respect to different level of drop-out

In the network reduction and reconstruction step, connected reactions were merged to form one metabolic module. The neural network model enables a non-linear dependency between gene expression and module flux. Hence, the flux rate could be determined by an “OR” operation of the high expression of any gene involved in the module, i.e., scFEA utilizes neighboring genes on the metabolic map to infer the metabolic flux of connected metabolic reactions, which increases robustness to dropout events and prediction accuracy. To further examine the method’s robustness, we simulated different levels of additional dropout events to our pancreatic cancer cell line data. Our data was collected by using the Smart-seq2-fluidigm protocol, whose original ratio of zero expressions of the metabolic genes is 50.22%. We simulated additional drop-out rate ranging from 4.34% to 34.78%, to reach a typical drop-out level of a droplet based scRNA-seq data (∼85%), and applied scFEA (see details in Supplementary Methods). We observed the total loss slightly increases from 0.1649 to 0.2722 when the zero ratio increased from 50% to 85% (**Figure 5G**). The module-wise and cell-wise correlation between the flux estimated from the original data and the perturbed data are consistently higher than 0.7437 and 0.8505 (**Figure 5H**), suggesting the high robust of scFEA with respect to different level of drop-out events.

### Application of scFEA on scRNA-seq data of tumor microenvironment revealed distinct metabolic stress, exchange and varied metabolic states in cancer and stromal cells

In this section, we majorly focused on validating the computational concept and applicability of scFEA on five real-world datasets, including two scRNA-seq data of cancer microenvironment, one single nuclei RNA-seq data of brain tissue, and one spatial transcriptomics data of breast cancer tissue. The data information is detailed in Supplementary Methods. All 169 metabolic modules across the whole metabolomic network was utilized in the analysis. Due to the lack of matched metabolomics information, we focused on demonstrating the capability of scFEA in inferring metabolic flux, metabolic stress, and cell groups and metabolic modules having distinct variations on these data sets.

#### Application on scRNA-seq data of cancer microenvironment

We applied scFEA on two publicly available scRNA-seq datasets collected from the microenvironment of melanoma (GSE72056) and head and neck cancer (GSE103322). In both data sets, we generated UMP based cell and cell group visualization by using predicted fluxomes of the 169 modules (**Figure 6A-D**). Interestingly, we identified that the metabolic flux distributions are quite homogeneous within cancer cells, but very distinctive comparing to immune and stromal cells in both data sets (**Figure 6A, C**). Distinct cell clusters of immune and stromal cells correspond to varied metabolic fluxomes were also identified (**Figure 6B, D**). A possible explanation is that cancer cells having a reprogrammed metabolism are more robust to the biochemical variations than immune and stromal cells in the tumor microenvironment.

**Figure 6.**
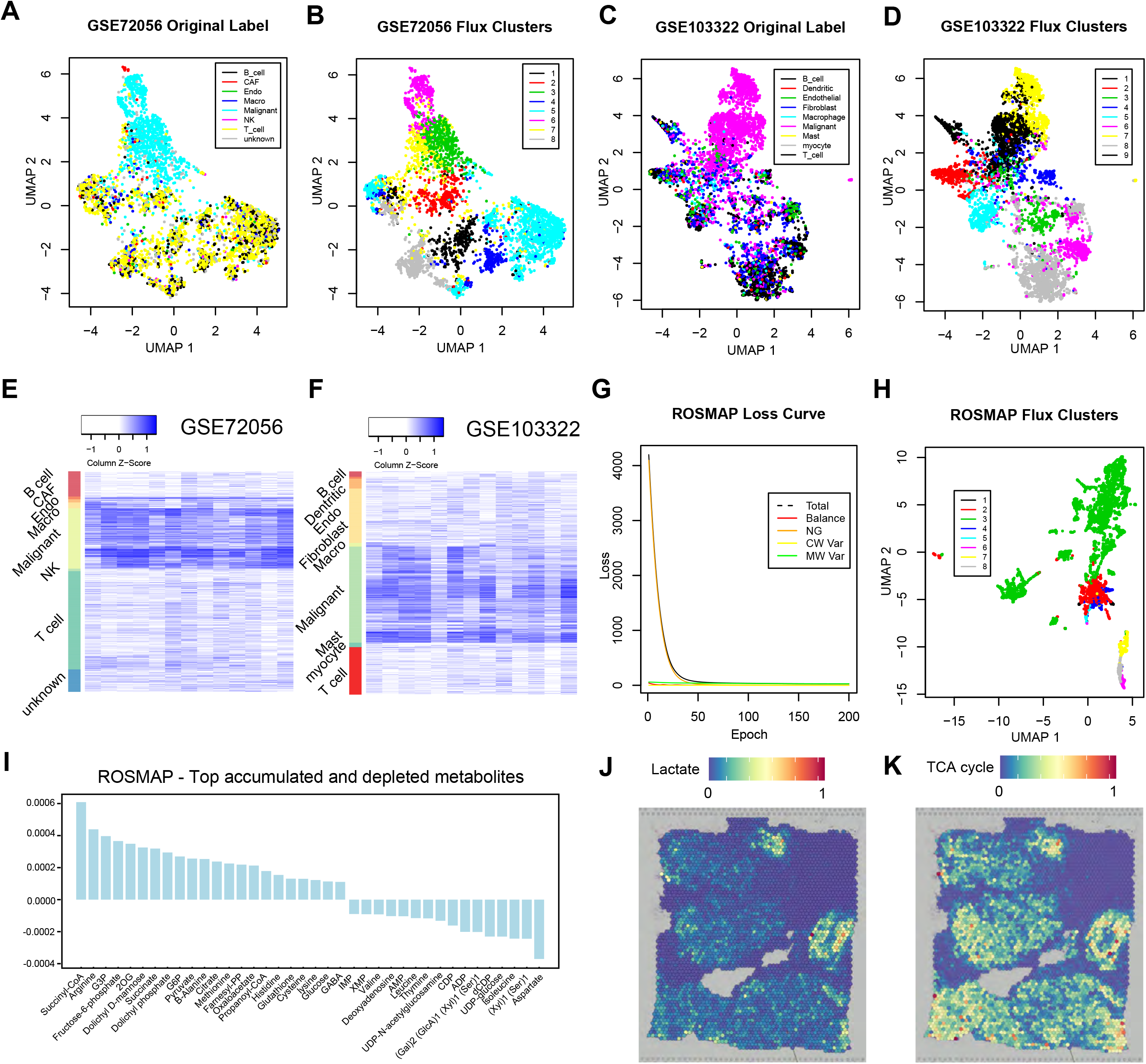
Application on two tumor scRNA-Seq datasets, ROSMAP, and one breast cancer spatial transcr iptomics dataset. (A) UMAP of the cell clustering based on metabolic fluxes of the GSE72056 melanoma data, the cell label was provided in original work. (B) UMAP of the cell clustering based on metabolic fluxes of the GSE72056, k-means clustering was used for cell clustering. (C) UMAP of the cell clustering based on metabolic fluxes of the GSE103322 head and neck cancer data, the cell label was provided in original work. (D) UMAP of the cell clustering based on metabolic fluxes of the GSE103322, k-means clustering was used for cell clustering. (E) Distribution of predicted cell-wise flux of glycolytic and TCA cycle modules of GSE72056 melanoma data. Each row is one cell, where row side color bar represents 8 cell types. Each column is one module. The left five columns are glycolytic modules from glucose to acetyl-CoA, the 6^th^ column is the reaction from acetyl-CoA to Citrate, the 7^th^ column is the reaction from pyruvate to lactate, and the right six columns (8-13 columns) are TCA cycle modules from citrate to oxaloacetic acid. (F) Distribution of predicted cell-wise flux of glycolytic and TCA cycle modules of GSE103322 head and neck cancer data. Each row is one cell, where row side color bar represents 9 cell types, respectively. The column is same as (E). (G) UMAP of the cell clustering based on metabolic fluxes of the ROSMAP data. k-means clustering was used for cell clustering. (H) Convergency curve of the total loss and four loss terms during the training of neural networks on the ROSMAP data. (I) Top accumulated and depleted metabolites predicted in the AD neuron cells in the ROSMAP data. The y-axis is metabolism stress level (or level of accumulation and depletion), where a positive value represents accumulation while a negative value represents depletion. The x-axis are metabolites in a decreasing order of the accumulation level. (J) scFEA predicted flux rate of lactate product on the spatial breast cancer data. The color of each point represents the spatial-wise predicted lactate product rate. The spatial plot is overlaid on the tissue slice image. (K) scFEA predicted flux rate of TCA cycle (citrate to 2OG) on the spatial breast cancer data.

We observed that the malignant cells have the highest metabolic rates in most metabolic reactions comparing to other cell types in both melanoma and head and neck cancer, especially for the glucose and amino acids metabolic modules (**Figure 6E, F**). On average, the estimated flux of TCA cycle and lactate production account for 43.4% and 52.5% of the glycolysis flux in head and neck cancer and 65.3% and 46.1% of the glycolysis flux (with additional carbon flow from other metabolites such as glutaminolysis) in melanoma, respectively, while the ratio of lactate production is much lower in other cell types. Our observation clearly suggested the existence of Warburg effect and metabolic shift in cancer cells, which is consistent to our previous findings of high lactate production in melanoma (Xu et al. 2012).

We identified that the malignant cells have the highest metabolic stress, which is defined as the total imbalance of intermediate substrates, followed by fibroblast and endothelial cells, and then immune cells. Similar to the pancreatic cancer cell line data, we identified that both cancer and stromal cells in both cancer types tend to have depleted glucose, G1P and G6P. In addition, cancer cells tend to suffer from a high depletion level of acetyl-coA. On the other hand, TCA cycle intermediates and amino acids tend to be accumulated in cancer cells. These observations are consistent to the findings derived from quantitative metabolomics data collected on solid cancer (Hirayama et al. 2009).

Interestingly, we noticed that the direction of imbalance for most intermediate metabolites seem to be the same throughout different cell types, though the imbalance level is much lower in stromal cells comparing to cancer cells. A possible explanation is that these cells were collected in a small region of the same microenvironment, subject to similar microenvironmental stresses, such as hypoxia and altered pH level, which causes a similar impact on the metabolic landscape of cells of different types. The predicted cell type specific fluxome and imbalance level of metabolites were given in Supplementary Table S4.

#### Application on droplet based scRNA-seq data of Alzheimer’s disease

We also applied scFEA on the ROSMAP snRNA-seq data (single nuclei RNA sequencing) collected from cells in the central nervous systems of Alzheimer’s disease (AD) patients and healthy donors (Mathys et al. 2019). Specifically, the ROSMAP snRNA-seq data was collected by using the 10x Genomics Chromium droplet-based protocol. Comparing to the Smart-seq based scRNA-seq data, droplet based data often have lower total expression signals and higher dropout rate. scFEA has been successfully applied on this data set. Changes of the total loss over the running epochs suggested the total loss converge to a small value during the training of the scFEA model (**Figure 6G**). Specifically, the flux balance loss forms the major loss term in the beginning of the training and quickly converge to a small value, suggesting a solution of high flux balance has been successfully identified in this data set. We identified that metabolic activity in neuron cells is higher than in other brain cell types. Cell clusters of different metabolic states were computed (**Figure 6H**), in which a large cluster consisting of cells with more activated metabolism has been identified (green labeled). We further studied on the metabolic stress of this cell cluster, which is enriched by neuron cells from AD patients (**Figure 6I**). We found that glucose, glycolytic and TCA cycle substrates, and glutathione are among the top accumulated metabolites. Suppressed glycolysis and dysfunctional TCA cycle that may lead to increased glucose and other intermediate metabolites, and elevated glutathione in response to reactive oxygen species, have been reported in AD (Atamna and Frey II 2007; Mandal et al. 2019; Le Douce et al. 2020). On the other hand, molecules involve in DNA synthesis and valine/leucine/isoleucine metabolism are most depleted in the AD neuron cells, which are consistent to the recently reported observations of suppressed DNA synthesis and valine metabolism in AD (Yurov et al. 2011; Polis and Samson 2020). More interestingly, we predicted aspartate and metabolites involved in glycosaminoglycan synthesis are greatly depleted in the AD neuron cells. Previous studies reported the association of these metabolites to AD (Doraiswamy 2003) (Huynh et al. 2019), however, their abundance change has been less studied. We anticipate that the cell-wise metabolic stress prediction enabled by scFEA could offer novel and systematic insight for biomarker prioritization.

#### Application on spatial transcriptomics data

Distinct cell clusters of different metabolic states were identified in the cancer microenvironment data (GSE72056 and GSE103322). We speculate that the different metabolic states are caused by varied biochemical conditions, such as hypoxia or oxidative stress level, in the tumor microenvironment. To further validate this hypothesis and the method, we applied scFEA on a spatial transcriptomics data of human breast cancer collected from 10x genomics visium protocol. Clearly, cells that are spatially near each other should be exposed to similar biochemical stress conditions. We predicted spatial spot specific metabolic flux, by first applying scFEA on the spatial the gene expression profile and then conducting associations of the predicted flux with spatial positions. scFEA identified two distinct spatial regions of high lactate production flux (**Figure 6J**) and six spatial regions of high TCA cycle flux (**Figure 6K**). Ratio of pyruvate → lactate flux and pyruvate → TCA cycle flux were computed, and the two high lactate production regions were predicted as of high hypoxia level, which were further validated by the high expression level of HIF1A regulated genes in cells of these regions.

## DISCUSSION

Despite a plethora of knowledge we have gained on metabolic dysregulation for many disease types, there are still major gaps in our understanding of the integrated behavior and metabolic heterogeneity of cells in the context of tissue microenvironment. Essentially, the metabolic behavior can vary dramatically from cell to cell due to the high metabolic plasticity, driven by the need to cope with various dynamic metabolic stress. Large amount of transcriptomics data obtained by scRNA-seq has proven to be endowed with the potential to deliver information on a cell functioning state and its underlying phenotypic switches. Designing advanced computational tools to harness the power of large scale scRNA-Seq data for reliable prediction of cell-wise metabolic flux and states is crucial to bridge the technological gap of single cell metabolomics. Knowledge derived therefrom will substantially benefit our understanding of the metabolic regulation intrinsic to diseased cells, and on factors imposed upon the diseased cells by its microenvironment and shed light on potential therapeutic vulnerabilities.

In sight of this demand, we developed a novel computational concept and method, namely scFEA, to predict metabolic flux at single cell resolution from scRNA-seq data, and the ultimate goal is to accurately construct and portray a compendium of metabolic states for different cell types and tissue contexts, and their relevance to various disease phenotypes. To experimentally validate scFEA, we generated an scRNA-seq data of a patient derived pancreatic cancer cells under four conditions of perturbed oxygen level and metabolic regulators, and matched tissue level metabolomics data and qRT-PCR profiles of key metabolic genes. We validated that the variations of metabolic flux predicted by scFEA are highly consistent with the observed metabolomic changes under different conditions. The scFEA predicted fluxome suggested the accumulation of glycolytic metabolites and depletion of TCA cycle metabolites, caused by suppression of the glycolysis pathway and TCA cycle pathways in both normoxia and hypoxia conditions. We also applied scFEA on in-drop or droplet based scRNA-seq data and spatial transcriptomics data. Our analysis suggested that scFEA could robustly predict cell and cell group specific metabolic shift for the data generated from different protocols. Notably, the fluxome estimated by scFEA enables a series of downstream analysis including identification of cell or tissue level metabolic stress, sensitivity evaluation of enzymes to the metabolic flux, and inference of cell-tissue and cell-cell metabolic exchanges.

The scFEA model has the following advantages: (1) the model characterizes true biological flux by leveraging the metabolic networks, and it is generally applicable as it requires only the input of scRNA-seq data; (2) the number of constraints, i.e. the number of flux balance conditions multiplied by the single cell number, is larger than the number of parameters, avoiding possible overfitting; and (3) The neural network based flux estimation can well handle the non-linear dependency between enzymatic gene expression and reaction rates. Theoretically, the scFEA model could be extended to estimate activity level of functional modules in a general biological network such as signaling pathways. The expression level of a signaling path reflects its capacity and the signaling molecules can be viewed as intermediates.

The neural network based optimization framework of scFEA enables a seamless integration of metabolomics data, kinetic parameters, spatial information, or other prior knowledge of metabolic imbalance, to better characterize cell and tissue level metabolic shifts of the target system. Specifically, metabolomics data, kinetic parameters or other prior knowledge can be utilized to better design the first layer of the neural network in modeling the flux of each module. Spatial information can be utilized to preselect group of cells for training spatially dependent model. A potential future direction is to implement the current flux estimation analysis in spatial transcriptomics to infer (1) metabolic shifts specific to spatial patterns and (2) metabolic exchange between adjacent cells. This application to spatial transcriptomics data will be particularly interesting for cancer studies, to reveal how the metabolism and macromolecule biosynthesis in stromal cells such as cancer associated fibroblast affect the adjacent cancer cells.

scFEA seeks for a constrained optimization of flux balance, where each flux was modeled as a non-linear function of the relevant enzymatic gene expression levels. The flux of each module is currently constrained to be scaled to the cell-wise total metabolic activity, *TA*_*j*_, to avoid trivia solution. However, our analysis suggested one *TA*_*j*_ for each cell may lead to similar metabolic flux distribution for different cells. Although our current setting has been validated by our matched scRNA-seq and metabolomics data, applications on publicly available cancer data suggested a similar metabolic imbalance trend among different cell types. We speculate that setting *T A*_*mj*_ for each super module *m* in cell *j* may increase the flexibility of cell specific metabolic imbalance, but at the price of possible over-fitting. A more sensitive approach is to train a specific model for each pre-defined cell group. The biological rationale is that the neural network parameters contain the information of “kinetic parameters” that link gene expression with metabolic rate, which differ among distant cell types, or cells under different conditions. Hence it is rationale to assume cell type specific parameters.

Overall, scFEA can efficiently delineate the sophisticated metabolic flux and imbalance specific to certain cell groups. We anticipate it has the potential to decipher metabolomic heterogeneity, and teasing out the metabolomic susceptibility to certain drugs, and ultimately warrant novel mechanistic and therapeutic insights of a diseased biological system at an unprecedented resolution.

## METHODS

### Collection of human metabolic map

We consider the human metabolic network as composed of different reaction types including metabolism, transport (including uptake and export), and biosynthesis. As detailed in Results, the reconstructed network consists of 22 super module classes of 169 modules. All reactions related to metabolism were collected from the Kyoto Encyclopedia of Genes and Genomes database (KEGG) (61). In total, 11 metabolism related super modules were manually summarized, which is comprised of glycolysis, TCA cycle, pentose phosphate, fatty acids metabolism and synthesis, metabolism of amino acids namely serine, aspartate, beta-alanine, glutamate, leucine/valine/isoleucine and urea cycle, propionyl-CoA and spermidine metabolism (Cao et al. 2017). The 11 metabolism super modules contain 1388 reactions, 317 enzymes, which corresponds to 563 genes.

Transporters enable the trafficking of molecules in and out of cell membranes. We collected the human transporter proteins, their corresponding genes and metabolite substrates from the Transporter Classification Database (Lin et al. 2015; Bhutia et al. 2016). In total, 80 transporter genes, and 35 related metabolites were collected.

An essential part of metabolic map is the biosynthesis pathways. KEGG database and literature (Moffatt and Ashihara 2002; DeAngelis et al. 2013; Zhang et al. 2015a; Zhang et al. 2015b; Krasnova and Wong 2016; Zulueta et al. 2016; Lv et al. 2017; Sun et al. 2018; Gao and Edgar 2019; Sun et al. 2020a; Sun et al. 2020b) are the main information sources used for building biosynthesis modules. We collected 69 biosynthesis modules forming 10 super modules, namely biosynthesis of hyaluronic acid, glycogen, glycosaminoglycan, N-linked glycan, O-linked glycan, sialic acid, glycan, purine, pyrimidine, and steroid hormones. Overall, the biosynthesis modules include 459 genes of 269 enzymes catalyzing 869 reactions.

More details of the collection of human metabolic map and the statistics of mouse metabolic map were provided in Supplementary Methods.

### Selecting genes of significant expression

We applied our inhouse method, LTMG, to determine the expression status of each genes in each single cell. LTMG considers the multi-modality of the expression profile of each gene throughout all the single cells, by assuming that the gene’s expression follows a mixture of suppressed state and activated states, as represented by the following likelihood function (Wan et al. 2019a).

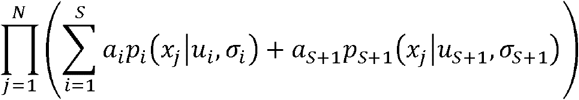

where *x*_*j*,_*j = 1*… *N* are the expression profile of gene *x* in *N* cell cells, the index 1 … *S* are the *S* active expression States and *S* + l is the suppressed expression state, *α*_*i*_ is the proportion of each state with *a*_l_ + … + *a*_*s*+1_ = 1, *a*_1…*S*_ > 0, *p*_*i*_,*u*_*i*_, and *σ*_*i*_ are the pdf, mean and standard deviation of each expression state. Specifically, LTMG considers the distribution of each mixing component, *p*_*i*_, as a left truncated Gaussian distribution, to account for the noise of drop out events. In this work, LTMG was used to fit to each gene’s expression and a gene’s expression and a gene was determined to have significant expression if 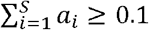, i.e., the gene has active expression states in at least 10% cells.

### Pre-filtering of active modules based on gene expression

Each metabolic module contains an input, an output, and a number of enzymes catalyzing the reactions. A reaction is considered as disconnected if none of the genes encoding its catalyzing enzymes is significantly expressed. A metabolic module is considered as blocked if there is no connected path from the input to the output. Considering the common drop-out events in scRNA-Seq data, especially for the drop-seq data, we adopted a conservative approach to pre-trim the metabolic modules: essentially, a module will be removed from further analysis if none of the genes involved in all reactions of this module has significantly active expressions.

### scFEA model setup and a belief propagation based solution of the flux model

#### Model Setup

We developed a novel optimization strategy to minimize L similar to the idea of belief propagation (Yedidia et al. 2001). Specifically, the flux balance of each metabolite 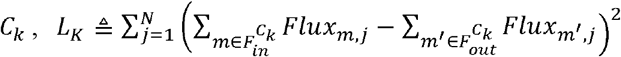, will be iteratively optimized, by taking into account all the Hop-2 neighbors in the factor graph (metabolites), denoted as *N*_*e*_(*C*_*k*_), and Hop-4 neighbors (metabolites), i.e., *N*_*e*_^2^(*C*_*k*_) := {*C*k_′_| *C*_*k*′_ ∈ *Ne*(*N*_*e*_(*C*_*k*_))\ *C*_*k*_}. Specifically, for a more efficient optimization, we adopt the idea of belief propagation by minimizing a reweighted flux imbalance: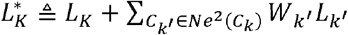 at each iteration, where *W*_*k*_ is a weight value in (0,1] representing the reliability of the current flux balance of *C*_*k* ′_. We set 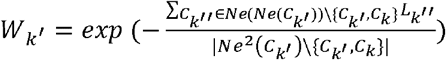 as an exponential function of the negative averaged imbalance level of 2-hop neighbors (metabolite) of *C*k_′_ excluding *C*_*k*_, with higher *W*_*k ′*_ denoting lower imbalance of the metabolites. The underlying idea is that the more reliable the current flux is estimated for the modules involving *C*_*k* ′_, which is reflected by the averaged imbalance level of its 2-hop neighbors, a higher weight *W*_*k ′*_> should be given to *C*_*k* ′_, such that when minimizing *LK*, a disruption of the flux balance of *C*_*k* ′_ will be more heavily penalized.

#### Neural network model setup

For each module, a neural network is used to represent the non-linear dependency between gene expressions and reaction rates. Each neural network has *a*1 hidden layers each with *a2* hidden nodes, and one output node. In this study, we took *a*1 = 3 and *a2 = 8*. A Hyperbolic Tangent activation function,*Tanhshrink* (*x*) = *x* – *tanh*, is used. The number of nodes and the number of hidden layers determines the complexity of network structure, which impacts the convergence time of optimization. Too simple structure would weaken the non-linear capability while too complex structure cause difficult to train all parameters and convergence. Our organized metabolic modules have an average gene number of 8, which determine the input nodes of scFEA. Since scFEA has 169 parallel subnetworks for each metabolic module, three hidden layers can leverage the level of non-linearity and overfitting and ensure a feasible computational cost (see details in Supplementary Methods).

#### Clustering analysis of cells with distinct metabolic states

scFEA adopts an attributed graph clustering approach to identify the group of cells and metabolic modules forming a distinct metabolic state. Three clustering approaches were provided to the results of scFEA for different tasks, namely clustering of (1) metabolic modules, (2) cells share a common state on the overall metabolic map, and (3) cells share a common state on selected metabolic modules.

#### Clustering of metabolic modules

Denote the adjacency matrix of the context specific metabolic map as *A*^*MxM*^ and predicted metabolic flux as *Flux*^*MxN*^, where *Fluxm,j* represents the predicted flux rate of the module in cell *j*, a two-stage spectral clustering was applied to cluster the metabolic modules based on and predicted *Flux*^*MxN*^. It is noteworthy here the *Flux*^*MxN*^ is usually much denser than the input scRNA-seq data since the metabolic modules modules without significant expression were excluded before the analysis. Specifically, denote, MxM as the *A*^*F,MxM*^ as the Euclidean distance of the *M* modules in *Flux*^*MxN*^, and *D*^*MxM*^ and *D*^*F,MxM*^ as the two diagonal matrices, in which 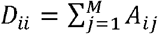 and 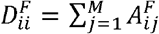. The normalized graph Laplacian matrices for the network topology and attributes similarity were defined as 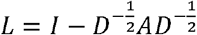 and 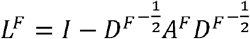.The normalized graph Laplacian matrices scale the topology and attributes similarity into the same scale. Denote *d* (*R*i,*R*j) and *d*^*F*^(*Ri,R*j)as the Euclidean the metabolic modules *Ri* and R*j* of the smallest *P*1 eigenvectors of and the smallest *P2* eigenvectors of *L*^F^, the modules were clusters by the *K-mean* method with using the following distance:

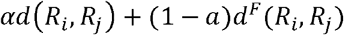

here *α, P*1 and *P*2, and the number of clusters are hyperparameters. Our empirical analysis suggested a default setting as *α*1 = 0.3, which assigns a higher weight to the similarity of the predict flux; 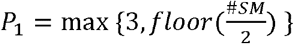 where *<u>#SM</u>* is the number super-modules in the current metabolic map; and 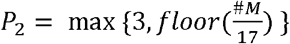, where *<u>#M</u>* is the number of non-zero modules in the current metabolic map. The number of clusters should be pre-given by users, which depends on the number of cells, cell types, and metabolic modules.

#### Clustering of cells

For a given metabolic map or a predefined group of metabolic modules, such an identified module cluster, scFEA conducts cell clustering analysis by using the spectral clustering approach based on the as defined above.

### Analysis of cell group specific metabolic stress and metabolic exchanges among cell groups

The cell-wise metabolic flux estimated by scFEA enables the analysis of metabolic stress. For a pre-defined cell group such as cells of the same type, the total imbalance of each compound will be computed and ranked. One-way t-test was applied to test if the imbalance is significantly different to 0. The metabolic exchange among different cell groups from one tissue sample were identified as the metabolites with different sign of metabolic imbalance in different cell groups, such as accumulation and depletion, or exporting or importing. Tissue level metabolic stress is computed as the total imbalance throughout multiple cells.

### Perturbation analysis

scFEA encodes a perturbation analysis to evaluate the impact of the change of each gene on the whole metabolic map. The perturbation analysis includes three components: (1) the direct impact of each gene 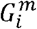 to the flux module *m* can be directly computed by its derivative 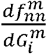 for all the modules containing 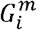 ; (2) the impact of the flux change of one module on other modules and flux balance of metabolites can be computed as the difference of flux of other modules estimated by scFEA while fixing the flux of module at different values; (3) the impact of each gene’s expression to the flux of distant modules and the flux balance was evaluated by integrating the approach of (1) and (2), i.e. first computing the flux change of the modules containing the gene and then evaluating the change of other modules and flux balance of other metabolites.

### Scalability analysis

The most time-consuming step in scFEA comes from the training of neural networks from each module to minimize the loss function. Maturing of a neural network consists of forward pass process and back-propagation update process. Forward pass process can be formed as m a trix multiplication, where input multiply the weights on the link and plus the bias. Then activation function has *O* (1) time complexity. The time complexity is *O* (*e* * *N* * (*i* * *h* + *h* * *m*)) for three layers forwarding and back-propagation update, where *i* is input layer node number, *h* is hidden layer node number, is output layer node number, *N* is the cell number, is number of iterations. In our work 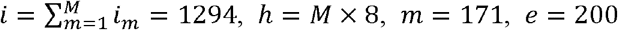 *h* = *M* × 8, *m* = 171, *e* = 200, *N* is cell number for each dataset. Paralleling GPUs matrix operation is encouraged since sub-networks are independent. We tested the time consumption of scFEA on a regular laptop of Intel i7-7600 CPU and 16GB RAM. The whole scFEA analysis process took 14 and 23 minutes on 4486 selected cells in the GSE72056 and 5902 selected cells in the GSE103322 datasets, respectively. Detailed scalability analysis is provided in Supplementary Methods.

### Patient-derived cell line models of pancreatic cancer

Pa03C cells were obtained from Dr. Anirban Maitra’s lab at The Johns Hopkins University (Jones et al. 2008). All cells were maintained at 37°C in 5% CO2 and grown in DMEM (Invitrogen; Carlsbad, CA) with 10% Serum (Hyclone; Logan, UT). Cell line identity was confirmed by DNA fingerprint analysis (IDEXX BioResearch, Columbia, MO) for species and baseline short-tandem repeat analysis testing in February 2017. All cell lines were 100% human, and a nine-marker short tandem repeat analysis is on file. They were also confirmed to be mycoplasma free.

### ScRNA-seq experiment

Cells were transfected with either Scrambled (SCR) (5′ CCAUGAGGUCAGCAUGGUCUG 3′, 5′ GACCAUGCUGACCUCAUGGAA 3′) or siAPEX1 (5′ GUCUGGUACGACUGGAGUACC 3′, 5′ UACUCCAGUCGUACCAGACCU 3′ siRNA). Briefly, 1×10^5^ cells are plated per well of a 6-well plate and allowed to attach overnight. The next day, Lipofectamine RNAiMAX reagent (Invitrogen, Carlsbad, CA) was used to transfect in the APEX1 and SCR siRNA at 20 nM following the manufacturer’s indicated protocol. Opti-MEM, siRNA, and Lipofectamine was left on the cells for 16 h and then regular DMEM media with 10% Serum was added.

Three days post-transfection, SCR/siAPEX1 cells were collected and loaded into 96-well microfluidic C1 Fluidigm array (Fluidigm, South San Francisco, CA, USA). All chambers were visually assessed and any chamber containing dead or multiple cells was excluded. The SMARTer system (Clontech, Mountain View, CA) was used to generate cDNA from captured single cells. The dscDNA quantity and quality was assessed using an Agilent Bioanalyzer (Agilent Technologies, Santa Clara, CA, USA) with the High Sensitivity DNA Chip. The Purdue Genomics Facility prepared libraries using a Nextera kit (Illumina, San Diego, CA). Unstrained 2×100 bp reads were sequenced using the HiSeq2500 on rapid run mode in one lane.

### ScRNA-seq data processing and analysis

FastQC was applied to evaluate the quality of the single cell RNA sequencing data. Counts were called for each cell sample by using STAR alignment pipeline against human GRCh38 reference genome. Cells with less than 250 or more than 10000 non-zero expressed genes were excluded from the analysis. Cells with more than 15% counts mapped to the mitochondrial genome were excluded as low quality cells, resulting 40 APEX1 KD and 48 Control cells under hypoxia condition and 27 APEX1 KD and 46 Control cells under normoxia condition for further analysis.

We utilized our in-house developed left truncated mixture Gaussian model to identify differentially expressed genes (Wan et al. 2019a). Pathway enrichment analysis of the genes in the identified bi-clusters are computed using hypergeometric test against the 1329 canonical pathway in MsigDB database (Liberzon et al. 2011), with p<0.001 as a significance cutoff.

### Metabolomic profiling and data analysis

To address the function of the mitochondria, S-1 Mitoplates (Biolog, Hayward, CA) *Mitochondrial Function Assay* were performed following the manufacturer’s protocol. The assay covers 14 metabolites in central metabolic pathways, namely glucose, glucose-1 phosphate, glucose-6 phosphate, pyruvate, and lactate in the glycolysis pathway, citrate, 2-oxoglutarate, succinate, fumarate, malate in the TCA cycle, and amino acids glutamate, glutamine, serine, and ornithine. Specifically, assay mix (60 minutes at 37°C) was added to the plates to dissolve the substrates. We collected, counted, resuspended PDAC cells in provided buffer and plated them at 5×104 cells/well after treatment (2020). Readings at 590nm were taken every 5 min for 4 hours at 37°C. Experiments were performed in triplicate with 3 biological replicates for the siAPEX1 and control PDAC cells under the normoxia condition. Raw data was analyzed using Graphpad Prism 8, and statistical significance was determined using the 2-way ANOVA and p-values <0.05 were considered statistically significant.

### qRT-PCR

qRT-PCR was used to measure the mRNA expression levels of the various genes identified from the scRNA-seq analysis. Following transfection, total RNA was extracted from cells using the Qiagen RNeasy Mini kit (Qiagen, Valencia, CA) according to the manufacturer’s instructions. First-strand cDNA was obtained from RNA using random hexamers and MultiScribe reverse transcriptase (Applied Biosystems, Foster City, CA). Quantitative PCR was performed using SYBR Green Real Time PCR master mix (Applied Biosystems, Foster City, CA) in a CFX96 Real Time detection system (Bio-Rad, Hercules, CA). The relative quantitative mRNA level was determined using the comparative Ct method using ribosomal protein L6 (RPL6) as the reference gene. Experiments were performed in triplicate for each sample. Statistical analysis performed using the 2−ΔΔCT method and analysis of covariance (ANCOVA) models, as previously published (Fishel et al. 2015).

Notation and abbreviation

| Abbreviation | Full name |
| --- | --- |
| **General** |  |
| scFEA | single-cell Flux Estimation Analysis |
| scRNA-seq | single cell RNA-sequencing |
| FBA | Flux Balance Analysis |
| ssGSEA | single sample gene set enrichment analysis |
| ANCOVA | analysis of covariance |
| PCC | Pearson Correlation Coefficient |
| KEGG | Kyoto Encyclopedia of Genes and Genomes |
| RELdb | Rate-Limiting Enzymes database |
| TFs | transcriptional factors |
| **Cell information** |  |
| Pa03c cell | pancreatic cancer cells |
| APEX1 KD | APEX1 knockdown |
| SCR | Scrambled |
| PV-ADSC | perivascular adipose tissue derived mesenchymal stem cells |
| MD | more differentiated |
| HS | high stemness |
| **Metabolites** |  |
| G1P | D-Glucose 1-phosphate |
| G6P | alpha-D-Glucose 6-phosphate |
| G3P | glyceraldehyde-3P |
| 3PD | 3-Phospho-D-glyceroyl phosphate |
| 2OG | 2-Oxoglutarate |
| OAA | oxaloacetate |

## DATA ACCESS

The raw and processed sequencing data of the normoxia scRNA-seq data generated in this study have been submitted to the NCBI Gene Expression Omnibus (GEO; https://www.ncbi.nlm.nih.gov/geo/)” www.ncbi.nlm.nih.gov/geo/) under accession number GSE99305. The scFEA package, full set of process scRNA-seq data, metabolomic profile, and analysis codes used in this work are available at https://github.com/changwn/scFEA.

## ACKNOWLEDGEMETS

This work was supported by NIH NIGMS 1R01GM131399-01, NSF IIS (N0.1850360), Showalter Young Investigator Award, and Precision Health Initiative of Indiana University. C.Z and S.C want to thank Dr. Ying Xu from the University of Georgia, Dr. Yunlong Liu, Dr. Kun Huang, Dr. Ji Zhang and Dr. Xiongbin Lu for their constructive suggestions and advices to this work.

## REFERENCES

2020. Mitochondrial Function Assays with MitoPlates. pp. https://www.biolog.com/products-portfolio-overview/mitochondrial-function-assays/.

Ahl PJ, Hopkins RA, Xiang WW, Au B, Kaliaperumal N, Fairhurst A-M, Connolly JE. 2020. Met-Flow, a strategy for single-cell metabolic analysis highlights dynamic changes in immune subpopulations. Communications Biology 3: 305.

Ali A, Abouleila Y, Shimizu Y, Hiyama E, Emara S, Mashaghi A, Hankemeier T. 2019. Single-cell metabolomics by mass spectrometry: Advances, challenges, and future applications. TrAC Trends in Analytical Chemistry.

Atamna H, Frey II WH. 2007. Mechanisms of mitochondrial dysfunction and energy deficiency in Alzheimer’s disease. Mitochondrion 7: 297–310.

Bhutia YD, Babu E, Ramachandran S, Yang S, Thangaraju M, Ganapathy V. 2016. SLC transporters as a novel class of tumour suppressors: identity, function and molecular mechanisms. Biochem J 473: 1113–1124.

Bishop AL, Rab FA, Sumner ER, Avery SV. 2007. Phenotypic heterogeneity can enhance rare-cell survival in ‘stress-sensitive’yeast populations. Molecular microbiology 63: 507–520.

Cao S, Zhu X, Zhang C, Qian H, Schuttler HB, Gong J, Xu Y. 2017. Competition between DNA Methylation, Nucleotide Synthesis, and Antioxidation in Cancer versus Normal Tissues. Cancer Res 77: 4185–4195.

Chen Y-P, Yin J-H, Li W-F, Li H-J, Chen D-P, Zhang C-J, Lv J-W, Wang Y-Q, Li X-M, Li J-Y. 2020. Single-cell transcriptomics reveals regulators underlying immune cell diversity and immune subtypes associated with prognosis in nasopharyngeal carcinoma. Cell Research 30: 1024–1042.

Damiani C, Maspero D, Di Filippo M, Colombo R, Pescini D, Graudenzi A, Westerhoff HV, Alberghina L, Vanoni M, Mauri G. 2019a. Integration of single-cell RNA-seq data into population models to characterize cancer metabolism. PLoS computational biology 15: e1006733.

Damiani C, Maspero D, Di Filippo M, Colombo R, Pescini D, Graudenzi A, Westerhoff HV, Alberghina L, Vanoni M, Mauri G. 2019b. Integration of single-cell RNA-seq data into population models to characterize cancer metabolism. PLoS computational biology 15: e1006733–e1006733.

DeAngelis PL, Liu J, Linhardt RJ. 2013. Chemoenzymatic synthesis of glycosaminoglycans: Re-creating, re-modeling and re-designing nature’s longest or most complex carbohydrate chains. Glycobiology 23: 764–777.

DeBerardinis RJ, Lum JJ, Hatzivassiliou G, Thompson CB. 2008. The biology of cancer: metabolic reprogramming fuels cell growth and proliferation. Cell metabolism 7: 11–20.

Doraiswamy PM. 2003. The role of the N-methyl-D-aspartate receptor in Alzheimer’s disease: therapeutic potential. Current neurology and neuroscience reports 3: 373–378.

Duncan KD, Fyrestam J, Lanekoff I. 2019. Advances in mass spectrometry based single-cell metabolomics. Analyst 144: 782–793.

Dunham I Kundaje A Aldred SF Collins PJ Davis C Doyle F Epstein CB Frietze S Harrow J Kaul R et al. 2012. An integrated encyclopedia of DNA elements in the human genome. Nature 489: 57–74.

Dunn L, Allen GF, Mamais A, Ling H, Li A, Duberley KE, Hargreaves IP, Pope S, Holton JL, Lees A. 2014. Dysregulation of glucose metabolism is an early event in sporadic Parkinson’s disease. Neurobiology of aging 35: 1111–1115.

Emara S, Amer S, Ali A, Abouleila Y, Oga A, Masujima T. 2017. Single-cell metabolomics. In Metabolomics: from fundamentals to clinical applications, pp. 323–343. Springer.

Evers TM, Hochane M, Tans SJ, Heeren RM, Semrau S, Nemes P, Mashaghi A. 2019a. Deciphering metabolic heterogeneity by single-cell analysis. ACS Publications.

Evers TMJ, Hochane M, Tans SJ, Heeren RMA, Semrau S, Nemes P, Mashaghi A. 2019b. Deciphering Metabolic Heterogeneity by Single-Cell Analysis. Analytical Chemistry 91: 13314–13323.

Feinberg AP. 2007. Phenotypic plasticity and the epigenetics of human disease. Nature 447: 433–440.

Fessenden M. 2016. Metabolomics: Small molecules, single cells. Nature 540: 153–155.

Fishel ML, Wu X, Devlin CM, Logsdon DP, Jiang Y, Luo M, He Y, Yu Z, Tong Y, Lipking KPJJoBC. 2015. Apurinic/apyrimidinic endonuclease/redox factor-1 (APE1/Ref-1) redox function negatively regulates NRF2. 290: 3057–3068.

Gao C, Edgar KJ. 2019. Efficient Synthesis of Glycosaminoglycan Analogs. Biomacromolecules 20: 608–617.

Gu W, Nowak WN, Xie Y, Le Bras A, Hu Y, Deng J, Issa Bhaloo S, Lu Y, Yuan H, Fidanis E. 2019. Single-cell RNA-sequencing and metabolomics analyses reveal the contribution of perivascular adipose tissue stem cells to vascular remodeling. Arteriosclerosis, thrombosis, and vascular biology 39: 2049–2066.

Hanahan D, Weinberg RA. 2011. Hallmarks of cancer: the next generation. cell 144: 646–674.

Harris RA, Wang T, Coarfa C, Nagarajan RP, Hong CB, Downey SL, Johnson BE, Fouse SD, Delaney A, Zhao YJ et al. 2010. Comparison of sequencing-based methods to profile DNA methylation and identification of monoallelic epigenetic modifications. Nat Biotechnol 28: 1097–U1194.

Heintzman ND, Stuart RK, Hon G, Fu YT, Ching CW, Hawkins RD, Barrera LO, Van Calcar S, Qu CX, Ching KA et al. 2007. Distinct and predictive chromatin signatures of transcriptional promoters and enhancers in the human genome. Nat Genet 39: 311–318.

Hirayama A, Kami K, Sugimoto M, Sugawara M, Toki N, Onozuka H, Kinoshita T, Saito N, Ochiai A, Tomita M et al. 2009. Quantitative Metabolome Profiling of Colon and Stomach Cancer Microenvironment by Capillary Electrophoresis Time-of-Flight Mass Spectrometry. Cancer Research 69: 4918–4925.

Hirschey MD, DeBerardinis RJ, Diehl AME, Drew JE, Frezza C, Green MF, Jones LW, Ko YH, L.A, Lea MA. 2015. Dysregulated metabolism contributes to oncogenesis. In Seminars in cancer biology, Vol 35, pp. S129–S150. Elsevier.

Honkoop H, de Bakker DE, Aharonov A, Kruse F, Shakked A, Nguyen PD, de Heus C, Garric L, Muraro MJ, Shoffner A. 2019. Single-cell analysis uncovers that metabolic reprogramming by ErbB2 signaling is essential for cardiomyocyte proliferation in the regenerating heart. Elife 8: e50163.

Huynh MB, Ouidja MO, Chantepie S, Carpentier G, Maïza A, Zhang G, Vilares J, Raisman-Vozari R, Papy-Garcia D. 2019. Glycosaminoglycans from Alzheimer’s disease hippocampus have altered capacities to bind and regulate growth factors activities and to bind tau. PloS one 14: e0209573.

Jaenisch R, Bird A. 2003. Epigenetic regulation of gene expression: how the genome integrates intrinsic and environmental signals. Nat Genet 33: 245–254.

Jones S, Zhang X, Parsons DW, Lin JC-H, Leary RJ, Angenendt P, Mankoo P, Carter H, Kamiyama H, Jimeno AJs. 2008. Core signaling pathways in human pancreatic cancers revealed by global genomic analyses.

Kanehisa M, Goto S. 2000. KEGG: kyoto encyclopedia of genes and genomes. Nucleic Acids Res 28: 27–30.

Kelley MR, Georgiadis MM, Fishel ML. 2012. APE1/Ref-1 role in redox signaling: translational applications of targeting the redox function of the DNA repair/redox protein APE1/Ref-1. Curr Mol Pharmacol 5: 36–53.

Kim J, DeBerardinis RJ. 2019. Mechanisms and Implications of Metabolic Heterogeneity in Cancer. Cell Metab 30: 434–446.

Kochanek KD, Murphy SL, Xu J, Arias E. 2019. Deaths: final data for 2017.

Krasnova L, Wong C-H. 2016. Understanding the Chemistry and Biology of Glycosylation with Glycan Synthesis. 85: 599–630.

Lan X, Roth S, Huttenlocher D, Black MJ. 2006. Efficient belief propagation with learned higher-order markov random fields. In European conference on computer vision, pp. 269–282. Springer.

Le Douce J, Maugard M, Veran J, Matos M, Jégo P, Vigneron P-A, Faivre E, Toussay X, Vandenberghe M, Balbastre Y. 2020. Impairment of glycolysis-derived L-serine production in astrocytes contributes to cognitive deficits in Alzheimer’s disease. Cell metabolism 31: 503-517. e508.

Lee D, Smallbone K, Dunn WB, Murabito E, Winder CL, Kell DB, Mendes P, Swainston N. 2012. Improving metabolic flux predictions using absolute gene expression data. BMC systems biology 6: 1–9.

Levine LS, Hiam KJ, Marquez DM, Tenvooren I, Contreras DC, Rathmell JC, Spitzer MH. 2020. Single-cell metabolic dynamics of early activated CD8 T cells during the primary immune response to infection. bioRxiv doi:10.1101/2020.01.21.911545:2020.2001.2021.911545.

Li X, Egervari G, Wang Y, Berger SL, Lu Z. 2018. Regulation of chromatin and gene expression by metabolic enzymes and metabolites. Nat Rev Mol Cell Biol 19: 563–578.

Liberzon A, Subramanian A, Pinchback R, Thorvaldsdóttir H, Tamayo P, Mesirov JPJB. 2011. Molecular signatures database (MSigDB) 3.0. 27: 1739–1740.

Lidstrom ME, Konopka MC. 2010. The role of physiological heterogeneity in microbial population behavior. Nature chemical biology 6: 705–712.

Lin L, Yee SW, Kim RB, Giacomini KM. 2015. SLC transporters as therapeutic targets: emerging opportunities. Nat Rev Drug Discov 14: 543–560.

Liu Y, Beyer A, Aebersold R. 2016. On the Dependency of Cellular Protein Levels on mRNA Abundance. Cell 165: 535–550.

Lv X, Cao H, Cao H, Lin B, Wang W, Zhang W, Duan Q, Tao Y, Liu X-W, Li X. 2017. Synthesis of Sialic Acids, Their Derivatives, and Analogs by Using a Whole-Cell Catalyst. Chemistry 23: 15143–15149.

Mandal PK, Shukla D, Tripathi M, Ersland L. 2019. Cognitive improvement with glutathione supplement in Alzheimer’s disease: A way forward. Journal of Alzheimer’s Disease 68: 531–535.

Mathys H, Davila-Velderrain J, Peng Z, Gao F, Mohammadi S, Young JZ, Menon M, He L, Abdurrob F, Jiang X. 2019. Single-cell transcriptomic analysis of Alzheimer’s disease. Nature 570: 332–337.

Matsuzawa Y. 2006. Therapy insight: adipocytokines in metabolic syndrome and related cardiovascular disease. Nature clinical practice Cardiovascular medicine 3: 35–42.

Mattson MP, Chan SL. 2001. Dysregulation of cellular calcium homeostasis in Alzheimer’s disease. Journal of Molecular Neuroscience 17: 205–224.

Mehrmohamadi M, Liu X, Shestov AA, Locasale JW. 2014. Characterization of the usage of the serine metabolic network in human cancer. Cell reports 9: 1507–1519.

Moffatt BA, Ashihara H. 2002. Purine and pyrimidine nucleotide synthesis and metabolism. Arabidopsis Book 1: e0018–e0018.

Polis B, Samson AO. 2020. Role of the metabolism of branched-chain amino acids in the development of Alzheimer’s disease and other metabolic disorders. Neural regeneration research 15: 1460.

Rask E, Olsson T, Soderberg S, Andrew R, Livingstone DE, Johnson O, Walker BR. 2001. Tissue-specific dysregulation of cortisol metabolism in human obesity. The Journal of clinical endocrinology & metabolism 86: 1418–1421.

Roadmap Epigenomics C, Kundaje A, Meuleman W, Ernst J, Bilenky M, Yen A, Heravi-Moussavi A, Kheradpour P, Zhang Z, Wang J et al. 2015. Integrative analysis of 111 reference human epigenomes. Nature 518: 317–330.

Robertson-Tessi M, Gillies RJ, Gatenby RA, Anderson AR. 2015. Impact of metabolic heterogeneity on tumor growth, invasion, and treatment outcomes. Cancer Res 75: 1567–1579.

Rohlenova K, Goveia J, García-Caballero M, Subramanian A, Kalucka J, Treps L, Falkenberg KD, de Rooij LP, Zheng Y, Lin L. 2020. Single-Cell RNA Sequencing Maps Endothelial Metabolic Plasticity in Pathological Angiogenesis. Cell Metabolism 31: 862-877. e814.

Saier MH, Jr., Tran CV, Barabote RD. 2006. TCDB: the Transporter Classification Database for membrane transport protein analyses and information. Nucleic Acids Res 34: D181–186.

Saurty-Seerunghen MS, Bellenger L, El-Habr EA, Delaunay V, Garnier D, Chneiweiss H, Antoniewski C, Morvan-Dubois G, Junier M-P. 2019. Capture at the single cell level of metabolic modules distinguishing aggressive and indolent glioblastoma cells. Acta Neuropathologica Communications 7: 1–16.

Schnell S. 2014. Validity of the Michaelis-Menten equation--steady-state or reactant stationary assumption: that is the question. FEBS J 281: 464–472.

Shah F, Goossens E, Atallah NM, Grimard M, Kelley MR, Fishel MLJMo. 2017. APE1/Ref-1 knockdown in pancreatic ductal adenocarcinoma–characterizing gene expression changes and identifying novel pathways using single-cell RNA sequencing. 11: 1711–1732.

Sun H, Zhang C, Cao S, Sheng T, Dong N, Xu Y. 2018. Fenton reactions drive nucleotide and ATP syntheses in cancer. J Mol Cell Biol 10: 448–459.

Sun H, Zhou Y, Skaro MF, Wu Y, Qu Z, Mao F, Zhao S, Xu Y. 2020a. Metabolic reprogramming in cancer is induced to increase proton production. Cancer Research 80: 1143–1155.

Sun H, Zhou Y, Skaro MF, Wu Y, Qu Z, Mao F, Zhao S, Xu Y. 2020b. Metabolic Reprogramming in Cancer Is Induced to Increase Proton Production. Cancer Res 80: 1143–1155.

Thompson C, Bauer D, Lum J, Hatzivassiliou G, Zong W-X, Zhao F, Ditsworth D, Buzzai M, Lindsten T. 2005. How do cancer cells acquire the fuel needed to support cell growth? In Cold Spring Harbor symposia on quantitative biology, Vol 70, pp. 357–362. Cold Spring Harbor Laboratory Press.

van der Knaap JA, Verrijzer CP. 2016. Undercover: gene control by metabolites and metabolic enzymes. Genes Dev 30: 2345–2369.

Vasdekis AE, Stephanopoulos G. 2015. Review of methods to probe single cell metabolism and bioenergetics. Metabolic engineering 27: 115–135.

Wagner A, Wang C, DeTomaso D, Avila-Pacheco J, Zaghouani S, Fessler J, Eyzaguirre S, Akama-Garren E, Pierce K, Ron-Harel N. 2020. In silico modeling of metabolic state in single Th17 cells reveals novel regulators of inflammation and autoimmunity. bioRxiv.

Wan C, Chang W, Zhang Y, Shah F, Lu X, Zang Y, Zhang A, Cao S, Fishel ML, Ma Q. 2019a. LTMG: a novel statistical modeling of transcriptional expression states in single-cell RNA-Seq data. Nucleic acids research 47: e111–e111.

Wan C, Chang W, Zhang Y, Shah F, Lu X, Zang Y, Zhang A, Cao S, Fishel ML, Ma Q et al. 2019b. LTMG: a novel statistical modeling of transcriptional expression states in single-cell RNA-Seq data. Nucleic Acids Res doi:10.1093/nar/gkz655.

Ward PS, Thompson CB. 2012. Metabolic reprogramming: a cancer hallmark even warburg did not anticipate. Cancer cell 21: 297–308.

Xiao Z, Dai Z, Locasale JW. 2019a. Metabolic landscape of the tumor microenvironment at single cell resolution. Nature communications 10: 1–12.

Xiao Z, Dai Z, Locasale JW. 2019b. Metabolic landscape of the tumor microenvironment at single cell resolution. Nature Communications 10: 3763.

Xiao Z, Locasale JW, Dai Z. 2020. Metabolism in the tumor microenvironment: insights from single-cell analysis. Oncoimmunology 9: 1726556.

Xu K, Mao X, Mehta M, Cui J, Zhang C, Xu Y. 2012. A comparative study of gene-expression data of basal cell carcinoma and melanoma reveals new insights about the two cancers. PloS one 7: e30750.

Yedidia JS, Freeman WT, Weiss Y. 2001. Generalized belief propagation. In Advances in neural information processing systems, pp. 689–695.

Yurov YB, Vorsanova SG, Iourov IY. 2011. The DNA replication stress hypothesis of Alzheimer’s disease. TheScientificWorldJOURNAL 11.

Zampieri M, Sekar K, Zamboni N, Sauer U. 2017. Frontiers of high-throughput metabolomics. Curr Opin Chem Biol 36: 15–23.

Zenobi R. 2013. Single-Cell Metabolomics: Analytical and Biological Perspectives. Science 342: 1243259.

Zhang C, Cao S, Toole BP, Xu Y. 2015a. Cancer may be a pathway to cell survival under persistent hypoxia and elevated ROS: a model for solid-cancer initiation and early development. Int J Cancer 136: 2001–2011.

Zhang C, Liu C, Cao S, Xu Y. 2015b. Elucidation of drivers of high-level production of lactates throughout a cancer development. J Mol Cell Biol 7: 267–279.

Zhang Y, Kim MS, Nguyen E, Taylor DM. 2020. Modeling metabolic variation with single-cell expression data. bioRxiv.

Zhao M, Chen X, Gao G, Tao L, Wei L. 2009. RLEdb: a database of rate-limiting enzymes and their regulation in human, rat, mouse, yeast and E. coli. Cell research 19: 793–795.

Zulueta MM, Lin SY, Hu YP, Hung SC. 2016. Synthesis of glycosaminoglycans. doi:10.1002/9781119006435.ch10, pp. 235–261.

