## Supplementary Figure S1 for "A graph neural network model to estimate cell-wise metabolic flux using single cell RNA-seq data"

### Glycolysis genes

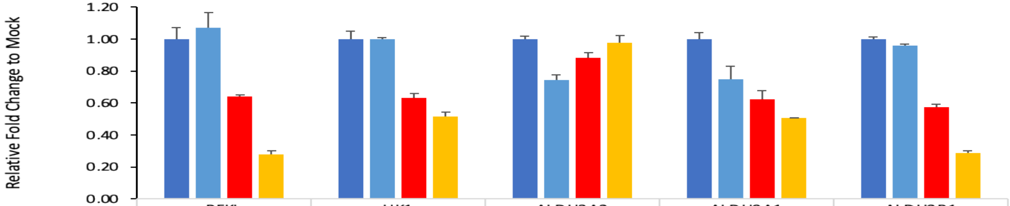


| | PFKL | HK1 | ALDH3A2 | ALDH9A1 | ALDH3B1 |
|---|---|---|---|---|---|
| Mock | 1.00 | 1.00 | 1.00 | 1.00 | 1.00 |
| SCR 30nM | 1.07 | 1.00 | 0.75 | 0.75 | 0.96 |
| siRef-1 10nM | 0.64 | 0.63 | 0.88 | 0.63 | 0.58 |
| siRef-1 30nM | 0.28 | 0.52 | 0.98 | 0.51 | 0.29 |

### TCA Cycle genes

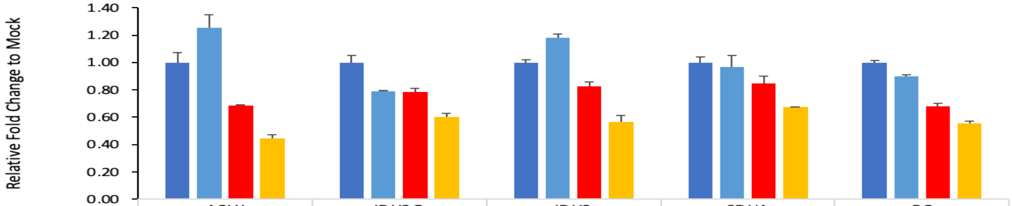


| | ACLY | IDH3G | IDH2 | SDHA | PC |
|---|---|---|---|---|---|
| Mock | 1.00 | 1.00 | 1.00 | 1.00 | 1.00 |
| SCR 30nM | 1.26 | 0.79 | 1.18 | 0.97 | 0.90 |
| siRef-1 10nM | 0.68 | 0.79 | 0.83 | 0.85 | 0.68 |
| siRef-1 30nM | 0.45 | 0.60 | 0.57 | 0.67 | 0.56 |

### OXPHOS genes

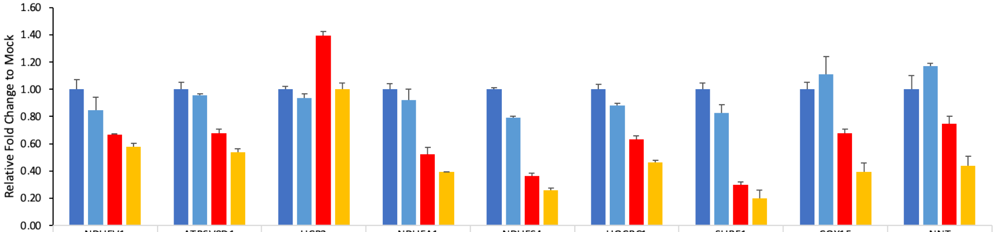


| | NDUFV1 | ATP6V0D1 | UCP2 | NDUFA1 | NDUFS4 | UQCRC1 | SURF1 | COX15 | NNT |
|---|---|---|---|---|---|---|---|---|---|
| Mock | 1.00 | 1.00 | 1.00 | 1.00 | 1.00 | 1.00 | 1.00 | 1.00 | 1.00 |
| SCR 30nM | 0.84 | 0.96 | 0.94 | 0.92 | 0.79 | 0.88 | 0.83 | 1.11 | 1.17 |
| siRef-1 10nM | 0.67 | 0.68 | 1.39 | 0.52 | 0.36 | 0.63 | 0.30 | 0.68 | 0.75 |
| siRef-1 30nM | 0.58 | 0.54 | 1.00 | 0.39 | 0.26 | 0.47 | 0.20 | 0.39 | 0.44 |
