## Supplementary figures and images for "A graph neural network model to estimate cell-wise metabolic flux using single cell RNA-seq data"

### Supplementary Figure S2

**Supplementary Figure S2**

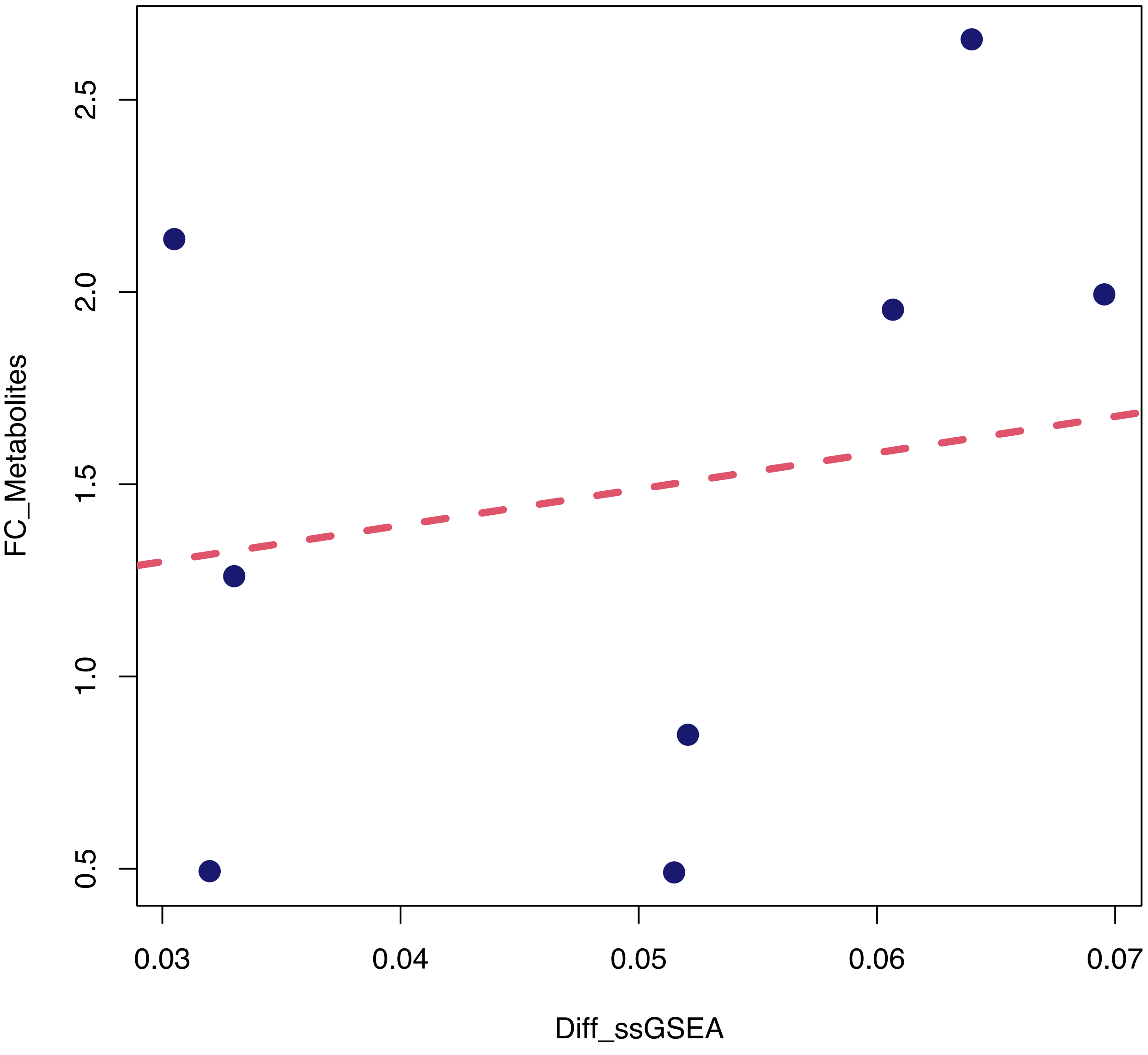

### Supplementary Figure S3

#### Supplementary Figure S3

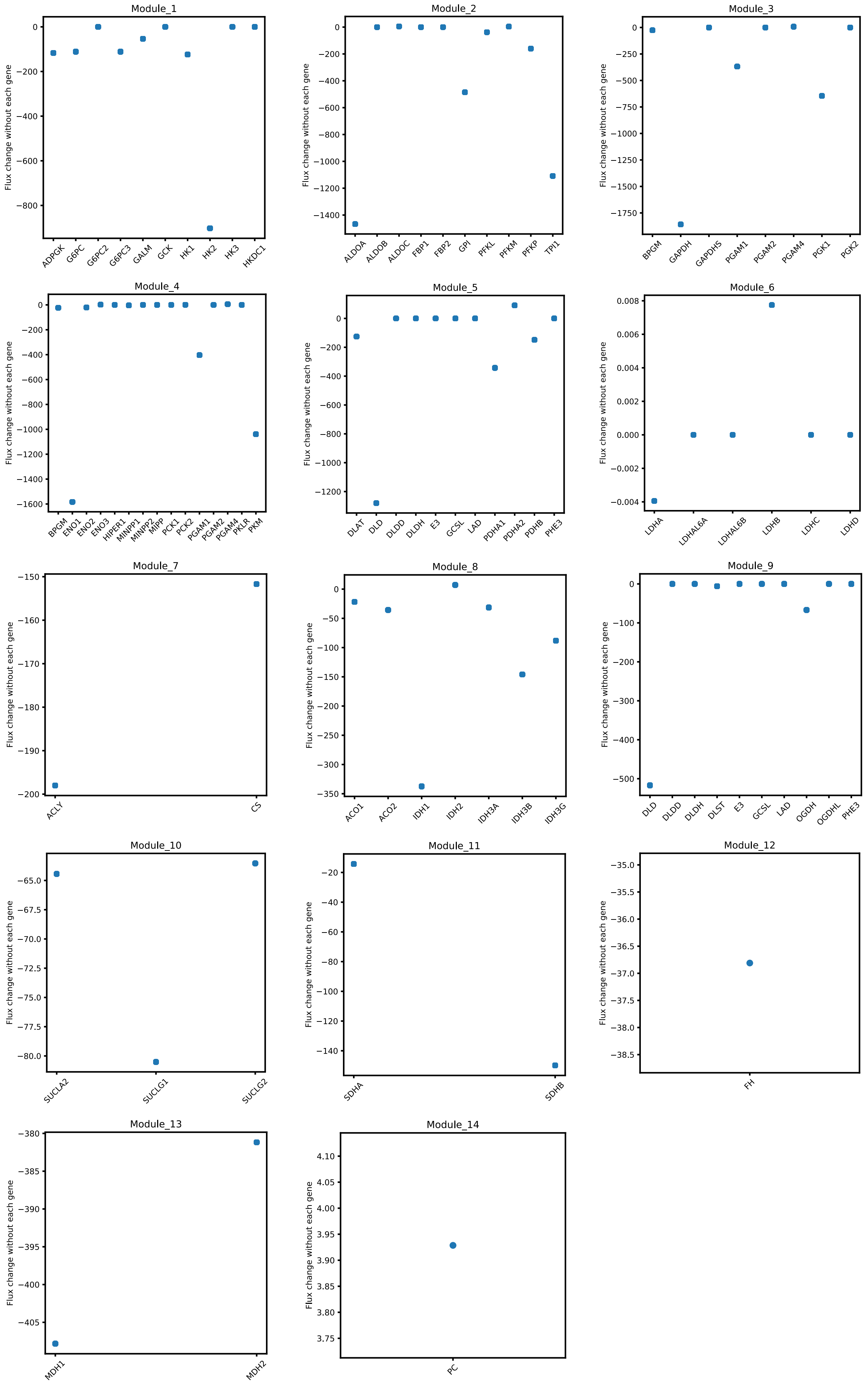

### Supplementary Figure S4

**Supplementary Figure S4**

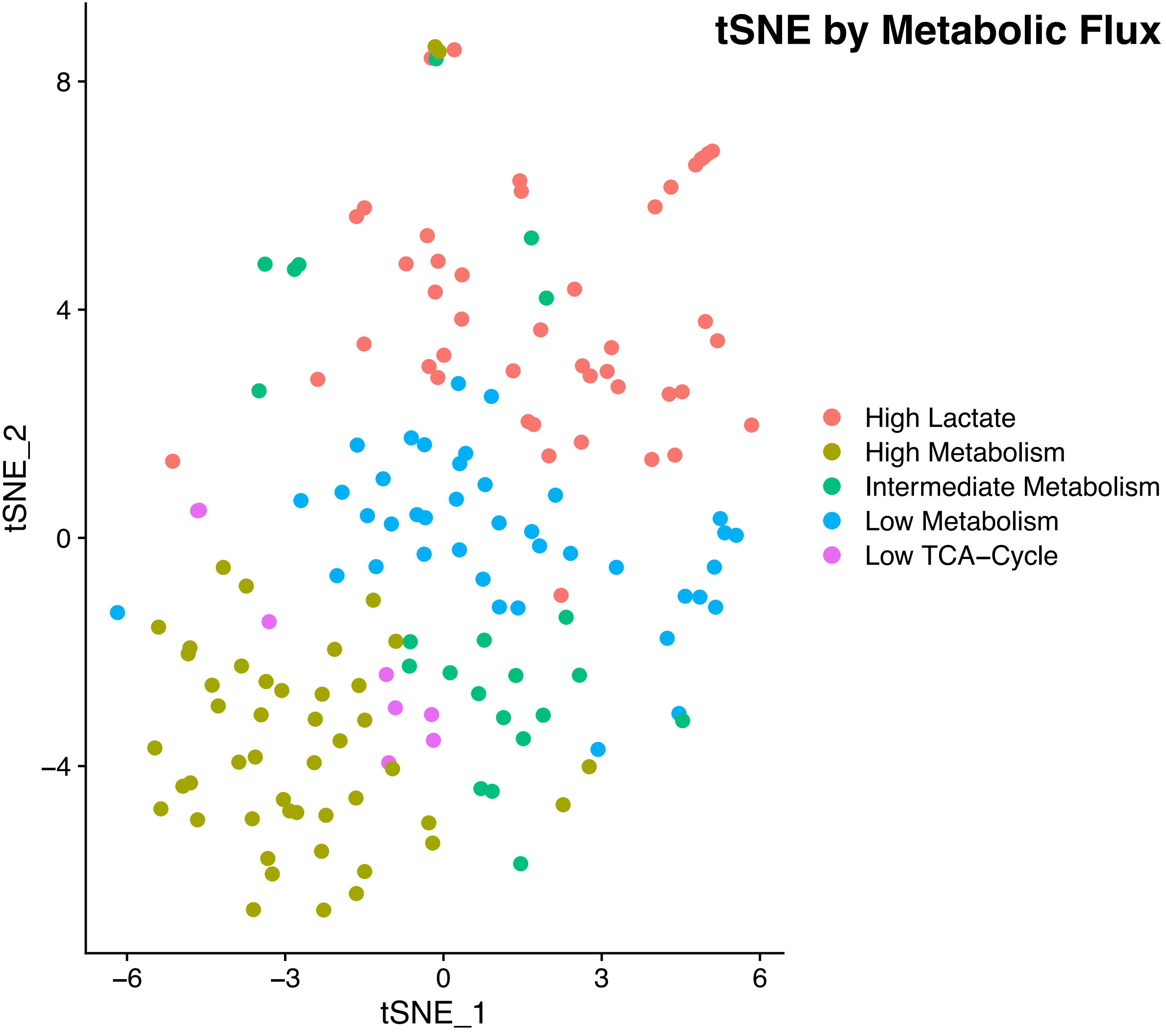

### Supplementary Figure S5

##### Supplementary Figure S5

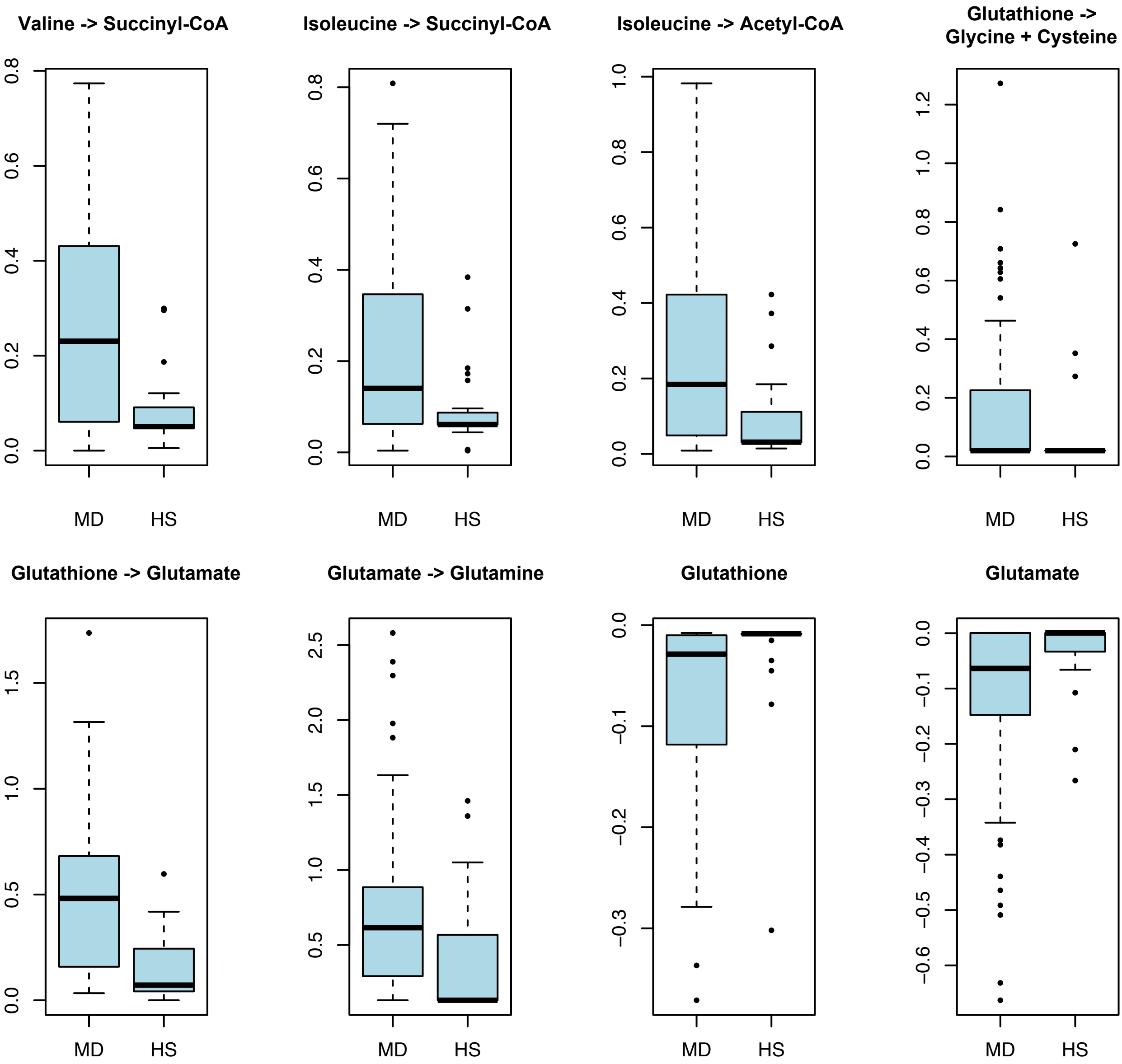

### Supplementary Figure S6

**Supplementary Figure S6**

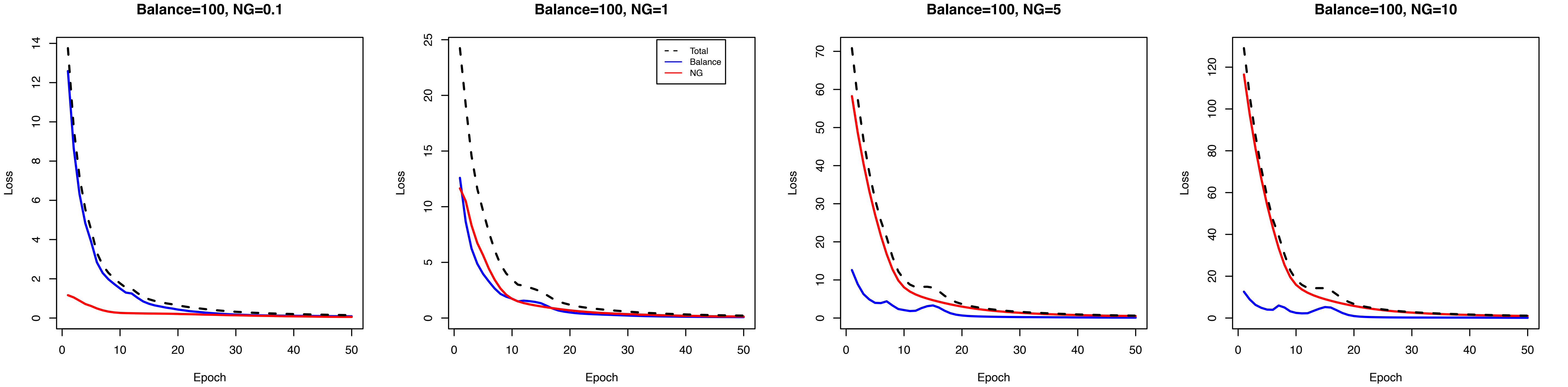
