## Supplementary Figure and Tables for "A graph neural network model to estimate cell-wise metabolic flux using single cell RNA-seq data"

**SUPPLEMENTARY FIGURES AND TABLES**

**Supplementary Figures**

**Supplementary Figure S1.** qRT-PCR results. Mock and SCR are controls and siRef-1 are knock down of APEX1.

**Supplementary Figure S2.** Correlation between metabolomic difference of the eight metabolites and differences of the averaged ssGSEA score of the modules using the eight metabolites as a substrate, in the APEX1-KD cells vs control. The x-axis is the difference of averaged ssGSEA score in the APEX1-KD cells vs control and the y-axis is the fold change of observed metabolomic profile.

**Supplementary Figure S3.** The impact of each gene to the metabolic module 1-14 (glycolysis and TCA cycle modules) in the Pa03c cell line data. The x-axis represents genes and y-axis represents impacts. The larger absolute value on the y-axis indicates a stronger impact of the gene to the metabolic module.

**Supplementary Figure S4.** tSNE plot of the cell clusters generated based on metabolic flux of the pancreatic cancer cell line data.

**Supplementary Figure S5.** Boxplots of the predicted fluxes of Valine -> Succinyl-CoA, Isoleucine -> Succinyl-CoA, Isoleucine -> Acetyl-CoA, Glutathione -> Glycine + Cysteine, Glutathione -> Glutamate, Glutamate -> Glutamine and predicted changes in the abundance of Glutathione and Glutamate in the PV-ADSC of high stemness (HS) and more differentiation (MD).

**Supplementary Figure S6.** Convergency of the flux balance loss and non-negative loss during the training of scFEA on the pancreatic cancer cell line data. The hyper parameters of the two loss were set differently to form four experiments. The flux balance loss, non-negative loss and total loss were blue, red and black-dash colored.

**Supplementary Tables**

Supplementary Table S1. Information of reorganized human metabolic map.

Supplementary Table S2. Differentially expressed genes (DEG) and Pathway Enrichment (PE) results of the Pa03c cell line data.

Supplementary Table S3. ssGSEA results, metabolomics data and clusters of metabolic modules derived in the Pa03c cell line data.

Supplementary Table S4. Predicted cell type specific fluxome and metabolic imbalance in the melanoma and head and cancer data.
